# How environmental drivers of spatial synchrony interact

**DOI:** 10.1101/2023.05.02.539141

**Authors:** Daniel C. Reuman, Max C.N. Castorani, Kyle C. Cavanaugh, Lawrence W. Sheppard, Jonathan A. Walter, Tom W. Bell

## Abstract

Spatial synchrony, the tendency for populations across space to show correlated fluctuations, is a fundamental feature of population dynamics, linked to central topics of ecology such as population cycling, extinction risk, and ecosystem stability. A common mechanism of spatial synchrony is the Moran effect, whereby spatially synchronized environmental signals drive population dynamics and hence induce population synchrony. After reviewing recent progress in understanding Moran effects, we here elaborate a general theory of how Moran effects of different environmental drivers acting on the same populations can interact, either synergistically or destructively, to produce either substantially more or markedly less population synchrony than would otherwise occur. We provide intuition for how this newly recognized mechanism works through theoretical case studies and application of our theory to California populations of giant kelp. We argue that Moran interactions should be common. Our theory and analysis explain an important new aspect of a fundamental feature of spatiotemporal population dynamics.

## 1 Introduction

Spatial synchrony, the tendency for geographically disjunct populations to show correlated fluctuations through time, is a fundamental feature of population dynamics, linked to important topics such as population cycling (Anderson *et al*., 2021), extinction risk (Ghosh *et al*., 2020c), and the stability of regional populations and ecosystem functioning (Wilcox *et al*., 2017). Though spatial synchrony (henceforth, *synchrony*) has been studied for decades in a wide variety of species ranging from viruses and plants to mammals, and at spatial scales ranging from centimeters to over 1000 km (Liebhold *et al*., 2004), recent advances in statistical methods and improvements in data availability have led to several major advances in our understanding of synchrony and its causes and consequences. For instance, the timescale structure of synchrony is now known to be important [Sheppard *et al*. (2016); Desharnais *et al*. (2018); Sheppard *et al*. (2019), see also Keitt (2008)], synchrony is now known to have a complex and pronounced geography (Defriez & Reuman, 2017a,b; Walter *et al*., 2017, 2020), and patterns of synchrony are now known to be asymmetric in distribution tails in a way that is important for system stability (Ghosh *et al*., 2020a,b; Walter *et al*., 2022). Our ability to infer the causes of synchrony is much improved in recent years (Sheppard *et al*., 2016; Defriez & Reuman, 2017b; Walter *et al*., 2017; Anderson *et al*., 2018; Sheppard *et al*., 2019), and there is also growing evidence that synchrony is changing as a newly recognized component of climate change (Defriez *et al*., 2016; Sheppard *et al*., 2016; Hansen *et al*., 2020). Distinct viewpoints on synchrony from population and community ecology are becoming integrated, leading to a more wholistic understanding of the importance of synchrony for ecosystem stability (Wang & Loreau, 2014). These new developments have led to an increasingly central role of the phenomenon of synchrony in many of the most important research topics in ecology.

Correlations between weather time series measured in different locations can induce synchrony between populations in those locations if the weather variables influence population processes. This mechanism, called the Moran effect, is now known to be one of the most important causes of synchrony. But the mechanism was originally proposed theoretically (Moran, 1953), and decades passed during which it was considered difficult to distinguish this potential cause of synchrony from others in field systems. Synchrony has long been thought to have three causes: dispersal, and trophic interactions with a synchronous or mobile species, in addition to the Moran effect (Liebhold *et al*., 2004). However, using common past statistical approaches which focussed on declines in population correlations with distance, patterns of synchrony produced by each of these mechanisms are quite similar (Ranta *et al*., 1999; Abbott, 2007; Liebhold *et al*., 2004; Walter *et al*., 2017); so examination of such patterns provides little or no traction for inferring the causes of synchrony. Early papers demonstrating the Moran effect mechanism resorted to special cases where dispersal was impossible and predators were absent (Grenfell *et al*., 1998; Tedesco *et al*., 2004). Controlled experiments also confirmed that all three theorized mechanisms could be involved in synchrony [e.g., Vasseur & Fox, 2009]. Nevertheless, the problem of inferring specific mechanisms of synchrony in field systems was considered a challenge until recently.

Recent research has provided new statistical viewpoints which have, when sufficient data are available, essentially solved the problem of inference of the causes of synchrony, and the research has revealed the broad importance of Moran effects. Approaches based on formal statistical comparisons between population synchrony maps and maps of environmental correlations produced inferences that precipitation and temperature Moran effects are important causes of synchrony of terrestrial (Defriez & Reuman, 2017b) and marine (Defriez & Reuman, 2017a) primary productivity. Another geographic approach, based on multiple regression of distance matrices (MRM), was used in an early paper to infer that a precipitation Moran effect is a cause of synchrony for the spongy moth (Haynes *et al*., 2013), an invasive forest pest whose name was recently changed from “gypsy moth.” Geographic approaches to identifying causes of synchrony, often MRM approaches specifically (Koenig *et al*., 2017; Walter *et al*., 2020; Bogdziewicz *et al*., 2021), have become widespread, and the geography of synchrony has become a mainstream area of study (Walter *et al*., 2017).

The approach of Gouveia *et al*. (2016), called “noncentered local indicators of spatial association” (ncLISA), is another geographic approach; as is the “fluvial synchrogram” of Larsen *et al*. (2021), a tool for studying synchrony in river networks. MRM approaches have also been used to identify dispersal as a cause of synchrony (Anderson *et al*., 2018), sometimes combined with genetic methods (Haynes & Walter, 2022). Another class of methods exploit the time and timescale structure of synchrony. Using a method that examined the match between temporal variations in population and environmental synchrony, Allstadt *et al*. (2015) confirmed the importance of a precipitation Moran effect for synchrony of spongy moth. A suite of wavelet and related Fourier techniques have been developed that can comprehensively describe the time and timescale structure of synchrony, identify Moran drivers of synchrony opperating in distinct timescale bands, and apportion fractions of observed synchrony to respective Moran drivers (Sheppard *et al*., 2016, 2017; Desharnais *et al*., 2018; Sheppard *et al*., 2019; Reuman *et al*., 2021; Anderson *et al*., 2021). The methods have been used, for instance, to provide one of the first clear demonstrations that changes in Moran effects are another of the impacts of climatic changes (Sheppard *et al*., 2016, 2019); and to illustrate the importance of the timescale structure of environmental covariation for altering Moran effects (Desharnais *et al*., 2018). These methods have since been used to identify Moran effects in several additional systems (e.g., Anderson *et al*., 2019; Walter *et al*., 2019, 2020; Garćıa-Carreras *et al*., 2022; Castorani *et al*., 2022).

The techniques reviewed above have made it possible for several recent papers to identify cases in which two or more distinct Moran drivers opperate simultaneously on the same populations (e.g., Defriez & Reuman, 2017b; Haynes *et al*., 2019; Walter *et al*., 2019, 2020; Anderson *et al*., 2021).

In fact, two recent papers have documented that Moran effects of distinct environmental drivers can interact, either synergistically or antagonistically, so that total population synchrony can be either greater than or less than what synchrony would be if the distinct Moran drivers operated independently (Sheppard *et al*., 2019; Castorani *et al*., 2022). Sheppard *et al*. (2019) showed that long-timescale (*>* 4y period) synchrony in a chlorophyll density index in the seas around the United Kingdom is substantially augmented by interactions between environmental drivers of synchrony and drivers linked with consumption by a copepod consumer. Castorani *et al*. (2022) showed that nutrient dynamics and wave disturbance, two Moran drivers of synchrony in giant kelp populations on the California (CA) coast, interact either synergistically or destructively, depending on which portion of the CA coast is examined and on the timescale of analysis.

The main purpose of this study is to establish a general theory of the mechanism of interacting Moran effects and to use examples and applications of the theory to provide ecological intuition for how the mechanism of interacting Moran effects works. Our goals are distinct from those of Kendall *et al*. (2000), who examined interactions between Moran and dispersal causes of synchrony. The basic fact that Moran drivers can interact, synergistically or destructively, can be demonstrated with a very simple model which we now elaborate, though understanding the full nature of the interaction mechanism will require the rest of this paper. Suppose *∈*^(1)^(*t*) and *∈*_*i*_^(2)^(*t*) are environmental random variables measured in locations *i* = 1, 2 at times *t*, and assume these are independent through time and standard normally distributed for all *i* and *t*. If a population index *w_i_*(*t*) follows the autoregressive process *w_i_*(*t*) = *bw_i_*(*t−*1)+*∈*^(1)^(*t*) for *i* = 1, 2, then the classic Moran theorem (Moran, 1953) implies that temporal population correlation equals temporal environmental correlation, i.e., cor(*w*_1_*, w*_2_) = cor(*∈*^(1)^, *∈*^(1)^). If, instead, *w_i_*(*t*) = *bw_i_*(*t −* 1) + *∈*^(1)^(*t*) + *∈*_*i*_^(2)^(*t*), so that both environmental variables influence populations, the Moran theorem then implies that population synchrony is cor(*w*_1_*, w*_2_) = cor(*∈*^(1)^ + *∈*^(2)^, *∈*^(1)^ + *∈*^(2)^). As we show in detail in SI section S1, it is straightforward to show that this quantity depends not only on the standard environmental synchrony measures cov(*∈*^(^*^a^*^)^, *∈*^(^*^a^*^)^), for *a* = 1, 2, but also on the “cross synchrony” measures cov(*∈*^(^*^a^*^)^, *∈*^(^*^b^*^)^) for *a /*= *b*. Cross synchrony measures represent synchrony between an environmental variable in one location and a different variable in another location. Fig. 1 illustrates, using this model, how population synchrony can therefore differ from what synchrony would be if the environmental processes *∈*^(1)^ and *∈*^(2)^ were unrelated, so that covariances between components of those processes are zero. The difference is due to interactions between the two Moran effects, and can be substantially positive or negative. This example was adapted from SI section S1 of Sheppard *et al*. (2019). All notation and abbreviations used throughout the paper are summarized in Table 1.

**Figure 1:**
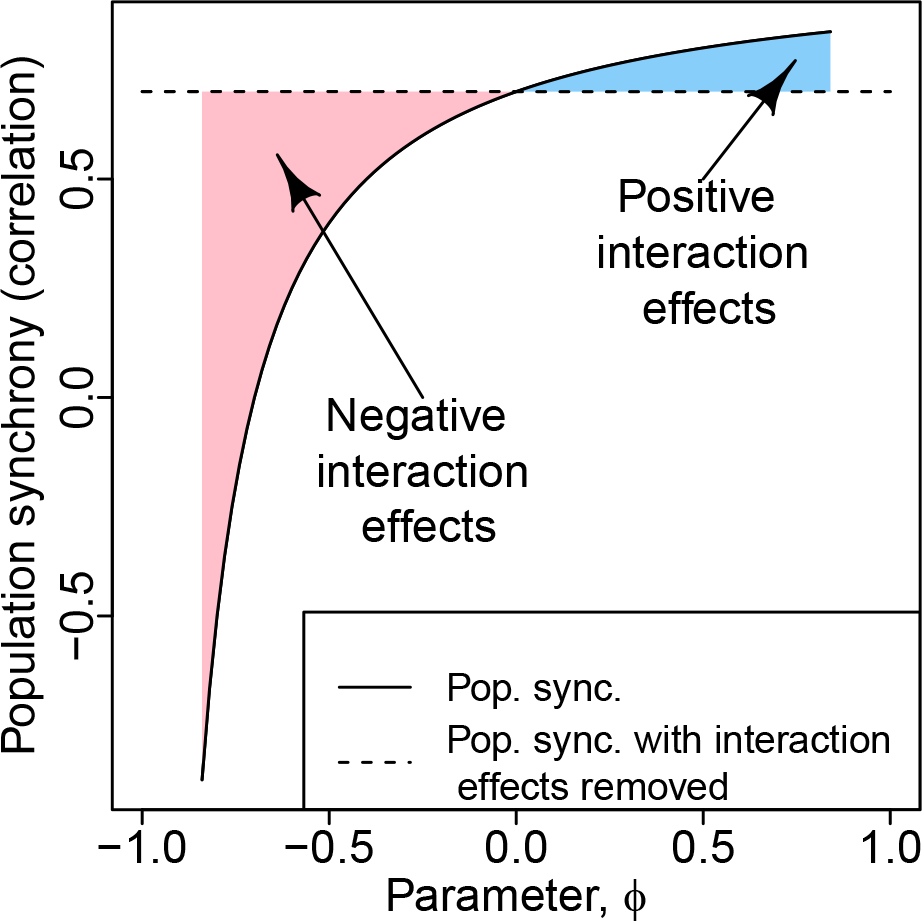
Example of interacting Moran effects. The example, which is based on the very simple model described in the Introduction, shows that interaction effects are possible, and can be substantially positive or negative. We used cov(*c*^(1)^*, c*^(1)^) = cov(*c*^(2)^*, c*^(2)^) = 0.7, and the values cov(*c*^(1)^*, c*^(2)^), for *i, j* = 1, 2, were all set equal to each other and to a parameter, *ϕ*, which appears on the horizontal axis and which characterizes the strength of cross synchrony of the environmental variables. See the online version for color renderings of all figures.

**Table 1:**
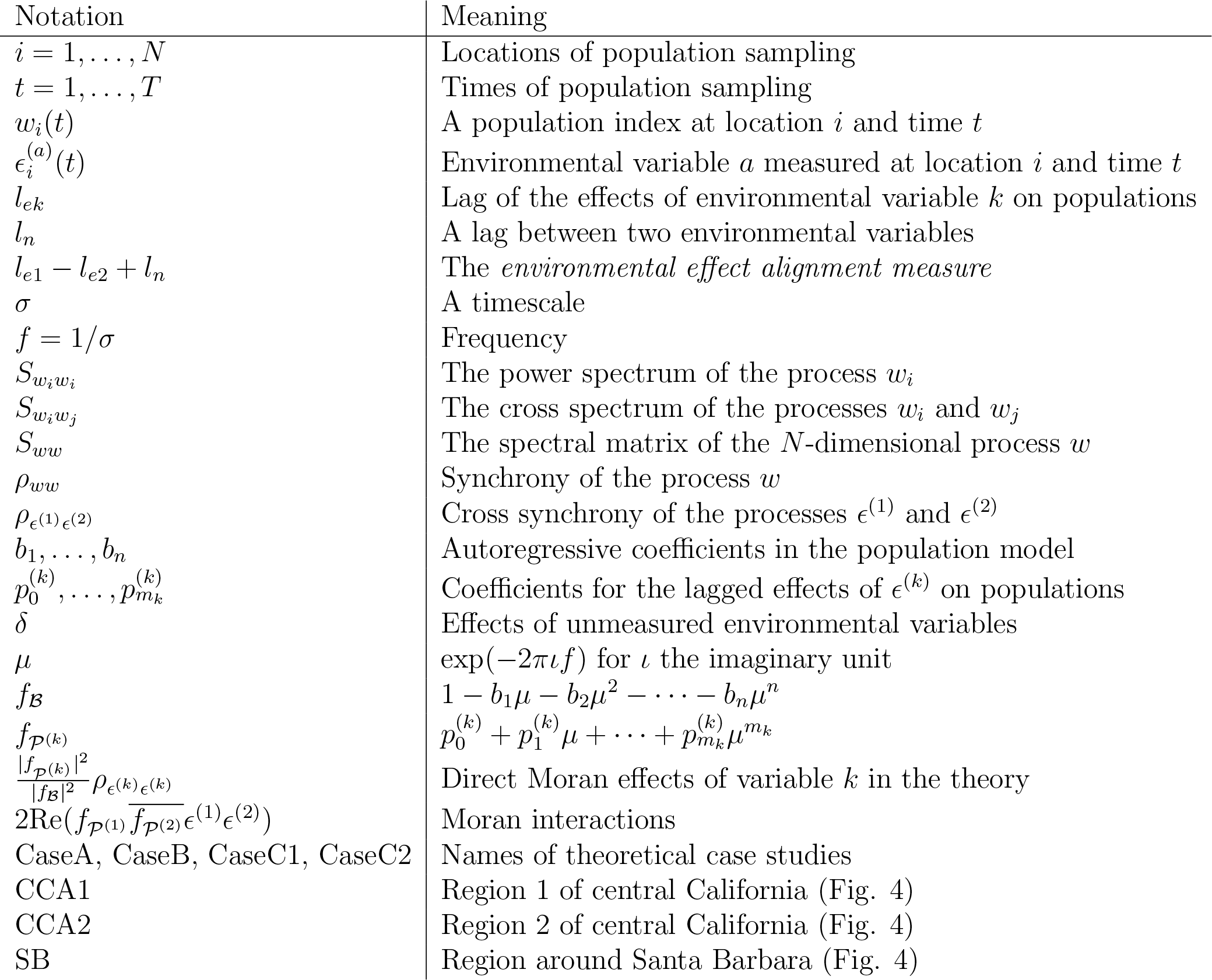
Summary of notation and abbreviations.

Having demonstrated that interactions between Moran effects can occur, we now use a simple analogy from common experience to begin to provide intuition. Consider *N* children, each riding on their own playground swing and each being pushed both by their own mother and their own father. Here, the children correspond to ecological populations and the parental pushing to environmental influences on the populations. If the fathers from seperate families were to synchronize their pushes, they would act as a Moran-like influence, tending to synchronize the swinging motions of the children. Likewise, if the mothers from seperate families were to synchronize their pushes, it would tend to have a separate synchronizing effect. If the pushes of the fathers were appropriately coordinated with the pushes of the mothers (either happening at the same time, if the mothers and fathers are standing on the same sides of the swinging children, or at opposite times in the swing period if the parents are on opposite sides), the children would swing higher, and would also swing more synchronously, demonstrating a synergistic interaction between synchronizing effects, i.e., a tendency for the two synchronizing effects to reinforce each other. On the other hand, if the maternal and paternal pushes were not appropriately coordinated, the children would swing more or less randomly, with smaller amplitude, and would become asynchronous with each other. This second case demonstrates an antagonistic interaction between synchronizing effects, i.e., a tendency for the two synchronizing effects to cancel. As oscillators, children on swings are distinct in many ways from populations, not least because swings oscillate on one frequency/timescale, whereas populations typically oscillate simultaneously on many timescales. We will see that a key insight is the use of a timescale-specific approach. By timescale-specific, we mean that fluctuations having different characteristic periods, which are superimposed in actual population time series data, can be understood separately. Timescale-specific approaches have been shown to be important to understanding synchrony and other phenomena (e.g., Sheppard *et al*., 2016; Anderson *et al*., 2021; Zhao *et al*., 2020). In the approach used in this paper, environmental and population signals are decomposed using Fourier analysis. The simple swing analogy turns out to supply, in much idealized form, the basic intuition our formal theory will extend.

Both Sheppard *et al*. (2019) and Castorani *et al*. (2022) argued that interactions between Moran effects may be a general feature of many systems, because of the large number of interrelated factors driving population fluctuations in most spatially extended systems. The same two studies demonstrated that interaction effects can be strong. For these reasons and others, development of a general theoretical and intutitive understanding of how interactions between Moran effects work is an important research goal. We explore in the Discussion why climate change may also influence Moran interactions, and we extend the arguments of Sheppard *et al*. (2019) and Castorani *et al*. (2022) that interactions between Moran effects are likely to be common.

Our specific goals for this study are as follows. First, (G1) we will elaborate a general theory of interacting Moran effects which allows a detailed understanding of how the mechanism works. Second, (G2) we will consider theoretical case studies, which emerge as important special cases of our general theory, and which illuminate the intuition behind how Moran effects may interact in real systems. Third, (G3) we will apply our theory to populations of giant kelp off the CA coast. Whereas Castorani *et al*. (2022) already carried out a detailed analysis of interacting Moran effects in CA kelp populations and their importance for kelp ecology, we instead use a simplified subset of the available kelp data. Our kelp analysis is intended to illuminate the inner workings of the general mechanism we describe rather than being an exploration of kelp dynamics, specifically. Overall, this study introduces a general theory of Moran interactions and uses it to conceptually illuminate this newly observed but potentially quite important mechanism of spatiotemporal population dynamics.

## 2 General Theory

### 2.1 Building intuition for Moran interactions: a single timescale

Prior to presenting our formal theory, we extend and formalize the intution behind it that began with the swing analogy in the Introduction. We again focus on a single timescale of oscillation, later combining timescales mathematically. Fig. 2a,b shows the period-20 Fourier components of two hypothetical, spatially synchronous environmental variables measured in sampling locations *i* = 1, 2, 3, and one way these can influence populations. The components are lagged, relative to each other, in the timing of their peaks (*l_n_* on the figure). They are also lagged in their effects on populations, by the amounts *l_e_*_1_ and *l_e_*_2_, respectively, i.e., peaks in the environmental signals manifest as maximum positive influence on populations after delays of *l_e_*_1_ and *l_e_*_2_, respectively. Such delays can be due to a variety of biological mechanisms associated with the life history of the organisms which comprise the populations. In this example, we assume for simplicity that larger values of both environmental variables are beneficial to populations, though the general theory described below does not require that assumption, and see also the next example which makes a different assumption. If, as in Fig. 2a, b, *l_e_*_1_ *− l_e_*_2_ + *l_n_* = 0 (or *l_e_*_1_ *− l_e_*_2_ + *l_n_* is any integer multiple of *σ* = 20, the period), then the periodic maximal positive influences of the two environmental variables coincide with each other as well as being spatial synchronous. This alignment of influences produces additional synchrony in populations, beyond what would manifest if environmental fluctuations were unrelated, because positive influences of variable 1 in location *i* will tend to coincide with positive influences of variable 2 in location *j*. In contrast, Fig. 2c, d shows the opposite scenario, for which *l_e_*_1_ *− l_e_*_2_ + *l_n_* = *−σ/*2. Thus the periodic maximal positive influence of environmental variable 1 coincides with the periodic maximal negative influence of environmental variable 2, reducing synchrony relative to the case of unrelated environmental variables. (The same outcome would occur if *l_e_*_1_ *−l_e_*_2_ +*l_n_* were any integer multiple of *σ* plus or minus *σ/*2.) Intermediate scenarios between the two scenarios of Fig. 2 are also possible, as will be revealed by the theory. We henceforth refer to the quantity *l_e_*_1_ *− l_e_*_2_ + *l_n_* as the *environmental effect alignment measure*, because it measures the extent to which the timing of the population influences of the two environmental variables are aligned. If we replace the assumption that larger values of both environmental variables are beneficial to populations with an assuption that larger values of the first variable and smaller values of the second are beneficial, scenarios of synergistic versus antagonistic Moran interactions are reversed, but with the same general concepts still operating (Fig. 3).

**Figure 2:**
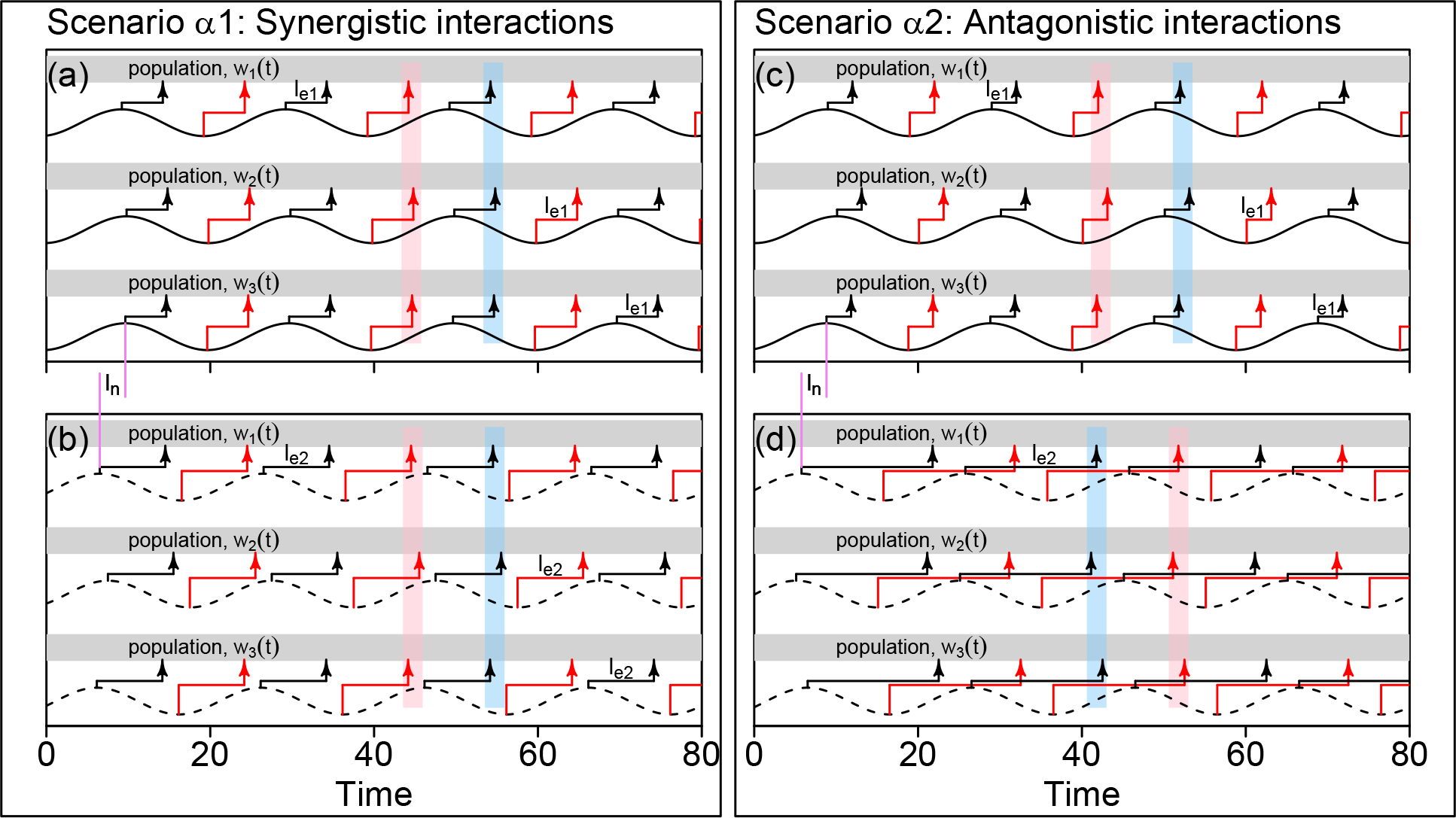
Illustration of the main concept of synergistically or antagonistically interacting Moran effects. Interactions require that each environmental variable itself be spatially synchronous; and then the alignment or misalignment of three types of lag determine the sign and strength of the interactions. Solid-line sinusoids represent the period-20 components of an environmental variable in three locations (*c*^(1)^ for *i* = 1, 2, 3) and dashed-line sinusoids represent the period-20 components of a different environmental variable in the same locations (*c*^(2)^ for *i* = 1, 2, 3). Black arrows represent peak positive influences of environmental variables on populations, which are lagged by an amount *l_e_*_1_ for *c*^(1)^ and by an amount *l_e_*_2_ for *c*^(2)^, where these lags differ across the scenarios *α*1 and *α*2, but are the same in all locations within one of these scenarios. Analogously, red arrows represent maximally negative effects. Due to environmental synchrony, peak positive population effects of the same variable occur at similar times across locations, and likewise for peak negative effects, illustrated with blue and pink rectangles. In the synergistic scenario (*α*1), the lag between the environmental variables (*l_n_*) and the lags of the population effects of the variables (*l_e_*_1_ and *l_e_*_2_) are aligned, i.e., the *environmental effect alignment measure*, *l_n_* + *l_e_*_1_ *− l_e_*_2_ (see main text), equals 0. So peak positive effects of *c*^(1)^ coincide with peak positive effects of *c*^(2)^ (the pink rectangles are aligned on a, b), augmenting synchrony. Likewise negative effects are aligned (blue rectan-gles). In the antagonistic scenario (*α*2), lags are misaligned, i.e., *l_n_* + *l_e_*_1_ *−l_e_*_2_ = *−σ/*2, where *σ* = 20 is the period. So peak positive effects of *c*^(1)^ coincide with maximally negative effects of *c*^(2)^, and maximally negative effects of *c*^(1)^ coincide with peak positive effects of *c*^(2)^ (pink rectangles on c are aligned with blue ones on d, and vice versa), reducing synchrony. See the online version for color renderings of all figures.

**Figure 3:**
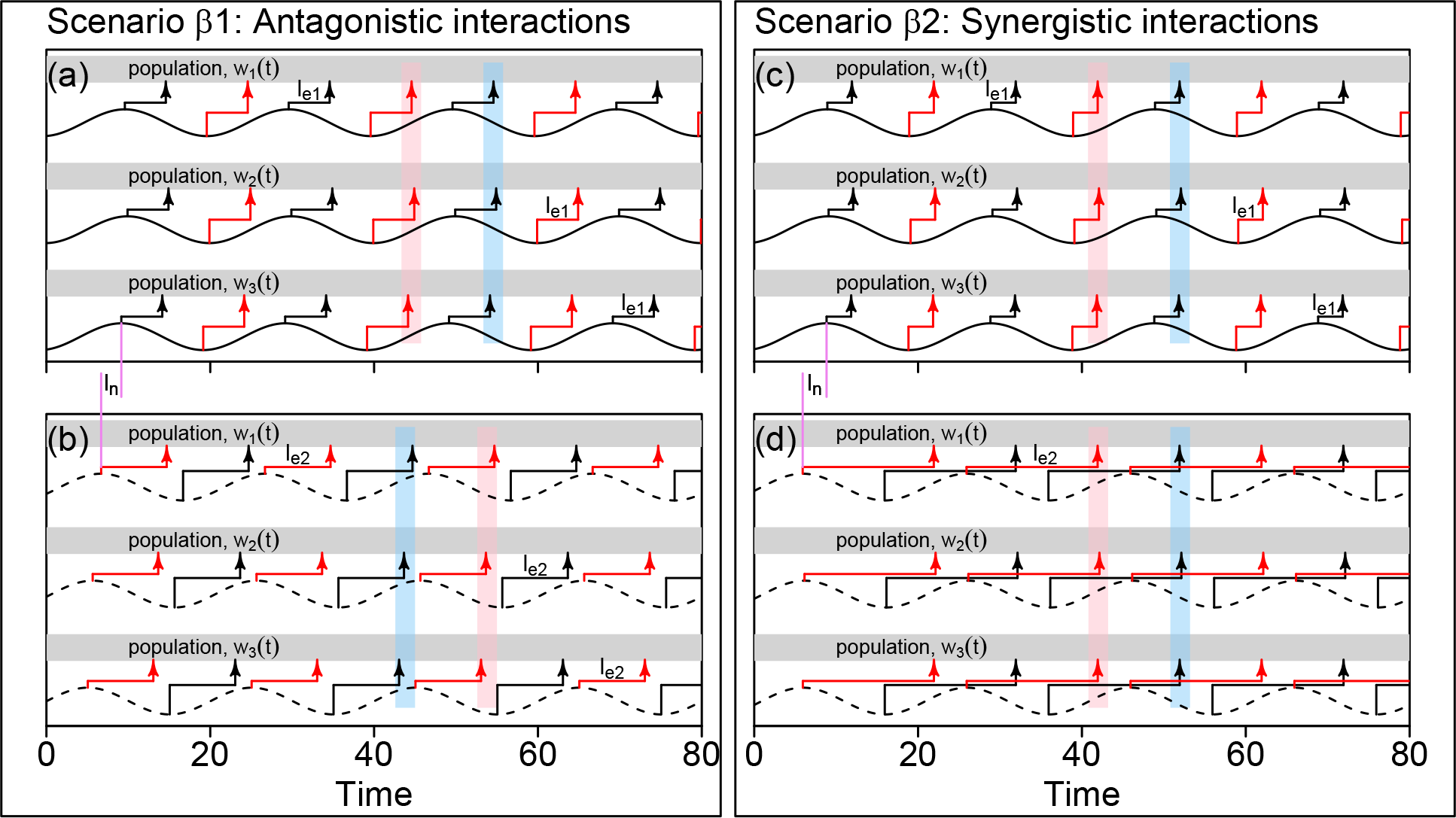
Similar to Fig. 2, but for slightly altered scenarios. For Fig. 2, for simplicity, we assumed that larger values of environmental variables had more positive influences on populations. We here assume that larger values of *c*^(1)^ have more positive population influences (as in Fig. 2), but that larger values of *c*^(2)^ have more negative population influences (contrasting with Fig. 2). In scenario *β*1, *l_n_* + *l_e_*_1_ = *l_e_*_2_, as in *α*1; but Moran interactions are nonetheless antagonistic because the negative effects of *c*^(2)^ means that the peak positive effects of that variable correspond to the peak negative effects of *c*^(1)^. In scenario *β*2, as in *α*2, the environmental effect alignment measure, *l_n_* + *l_e_*_1_ *− l_e_*_2_, equals *−σ/*2 for *σ* = 20 the period. But Moran interactions are synergistic because the negative effects of *c*^(2)^ means that the peak positive effects of that variable correspond to the peak positive effects of *c*^(1)^. See the online version for color renderings of all figures.

### 2.2 Formal theory

Our formal theory requires a conceptual understanding of the *spectrum* of an environmental or population time series, as well as of the *cospectrum* and *cross spectrum* of two time series, so we briefly introduce these concepts. If *w_i_*(*t*) is a stochastic process or time series measured at location *i* (e.g., the population density time series of a species of interest in that location), the spectrum *S_w__iwi_* (*f*) is a function of frequency, *f* . For a periodic ocillation, *f* can be defined as one over the timescale, or period, *σ*, of the oscillation. The spectrum *S_w__iwi_* (*f*) is larger for frequencies at which *w_i_*(*t*) oscillates more. So, for example, a population *w_i_*(*t*) that shows strong oscillatory dynamics of 5-year period will have a large value of *S_w w_* (*f*) for *f* = <u>^1^</u> y*^−^*^1^. The spectrum separates the overall variance of a time series into contributions which occur at different frequencies, in the sense that an integral of *S_w__iwi_* (*f*) across all frequencies equals var(*w_i_*). In a similar way, the cospectrum of two time series, *w_i_*(*t*) and *w_j_*(*t*), is a function of *f* that takes large values at frequencies for which oscillations in *w_i_*(*t*) and *w_j_*(*t*) are both strong and strongly correlated, i.e., they have the same or similar phase. Here, *j* refers to another location where measurements were taken. The cospectrum is the real part of the cross spectrum, *S_wiwj_*(*f*), which is a complex-valued function of frequency. The cross-spectrum takes large-magnitude values at frequencies, *f*, for which the oscillatory components of *w_i_*(*t*) and *w_j_*(*t*) are strong and in a consistent phase relationship to each other; and the complex phase of the cross spectrum at *f* then quantifies that relationship. The notation *S_ww_*(*f*) refers to the *spectral matrix*, which has *ij*^th^ entry *S_w w_* (*f*). Spectral methods are standard [e.g., Vasseur & Gaedke (2007); Defriez & Reuman (2017a)], and many background references are available [e.g., Brillinger (2001)].

Much prior work demonstrates the importance of a frequency- or timescale-specific approach to synchrony [e.g., Vasseur & Gaedke (2007); Keitt (2008); Defriez *et al*. (2016); Sheppard *et al*. (2016); Desharnais *et al*. (2018); Anderson *et al*. (2021); frequency- and timescale-specific approaches are equivalent because frequency and timescale are reciprocal], and it will turn out (see below, and Discussion) that a timescale-specific approach is essential to the development of our new theory.

We therefore here define, using the spectral methods outlined above, a frequency/timescale-specific measure of synchrony, as well as a new concept of *cross-variable synchrony*. If time series data *w_i_*(*t*) for *t* = 1*, … , T* were gathered at locations *i* = 1*, … , N*, our synchrony measure is simply 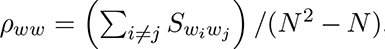, the average of the cross spectra for all pairs of distinct locations.

This is a real-valued function of frequency, an integral (across frequencies) of which is the classic, non-frequency-specific synchrony measure 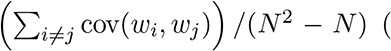 (see SI section S2 for details, here). If two time series *∈*^(1)^(*t*) and *∈*_*i*_^(2)^(*t*) (*t* = 1*, … , T* and *i* = 1*, … , N* ) were measured at each sampling location (e.g., two environmental variables), cross-variable synchrony (or, simply, *cross synchrony* ) between the variables is defined as 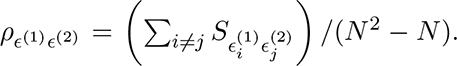. This is interpretable as pertaining to spatial synchrony because it makes comparisons across distinct locations, i.e., *i /*= *j*. It is interpretable as cross-variable synchrony because comparisons are between the two variables, i.e., cross spectra between time series components of *∈*^(1)^ and *∈*^(2)^ are used. The new index takes into account possible time lags. For instance, if both *∈*^(1)^ and *∈*^(2)^ show strong, spatially synchronous, 4-year-period oscillations, but peaks in the *∈*^(2)^ oscillations consistently lag peaks in the *∈*^(1)^ oscillations by a year, then *ρ_E_*(1)*_E_*(2) , which is complex valued, will have high magnitude at timescale 4y, equivalent to frequency 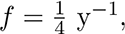, and will have phase at that frequency equal to *π/*2, reflecting the one-year lag. See SI section S3 for detailed examples.

Our population model is

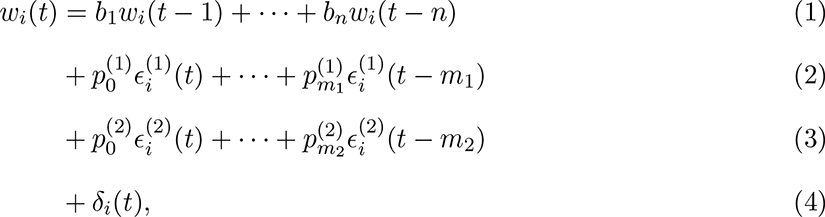

where *w_i_*(*t*) is an index of the population in location *i* = 1*, … , N* at time *t*, and *∈*^(1)^ = (*∈*^(1)^*, … , E*^(1)^), *∈*^(2)^ = (*∈*^(2)^*, … , E*^(2)^) and *δ* = (*δ*_1_*, … , δ_N_*) are environmental processes at the same locations. The processes *∈*^(1)^ and *∈*^(2)^ are taken to be measured, whereas *δ* represents the aggregate influence of unmeasured processes. We assume that the combined process (*∈*^(1)^*, … , E*^(1)^, *∈*^(2)^*, … , E*^(2)^*, δ*_1_*, … , δ_N_*) is an ergodic second-order stationary stochastic process (Brillinger, 2001) with expected values of its components equal to 0. We make additional mild regularity assumptions for model stability (SI section S4). This model can be seen as a linearization of a very general dynamical model, influenced by “weak noise” [see, e.g., SI section S1.2 of Desharnais *et al*. (2018) and SI section S1 of Walter *et al*. (2017)]. Linearization and “weak noise” assumptions have been commonly adopted to make theoretical progress in ecology, and it has been demonstrated that results based on a weak noise assumption often hold for noise which is fairly strong (Nisbet *et al*., 1977; Desharnais *et al*., 2018). See Brillinger (2001) for background on stochastic processes.

Fulfilling goal G1 of the Introduction (i.e., to elaborate theory of interacting Moran effects), the outcome of our theory is an equation which expresses population synchrony in terms of synchrony of the environmenal processes *∈*^(1)^ and *∈*^(2)^, and cross synchrony between those processes:

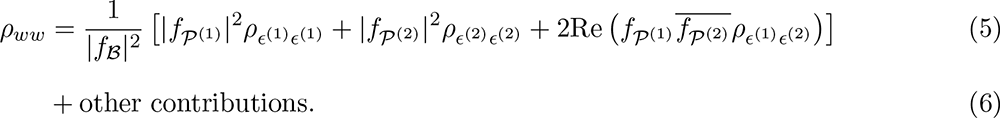

Here, 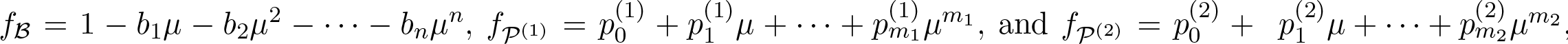, where *µ* = exp(*−*2*πιf*), and where *ι*, the Greek letter iota, is the imaginary unit. The derivation of the theory is in SI section S5.

Comparing the terms on the right of (5) gives the relative contributions of direct Moran effects and interactions between Moran effects. The first term on the right of (5) is the component of population synchrony due to the direct Moran effects of *∈*^(1)^, the second term is the component due to the direct Moran effects of *∈*^(2)^, and the third term is the component due to interactions between the Moran effects of the two drivers. The “other contributions” above correspond to synchronizing influences of *δ* and of interactions between *δ* and the *∈*^(^*^i^*^)^. Such contributions would be difficult to assess because *δ* was unmeasured.

The way direct Moran effects in our theory are interpreted is fairly straightforward. The magnitudes of the quantities *f_P_*(1) and *f_P_*(2) quantify the strength of influence of *∈*^(1)^ and *∈*^(2)^ on populations at the timescale *σ* = 1*/f* . The direct Moran effect term in (5) for *∈*^(1)^, i.e., *|f_P_*(1) |^2^*ρ_E_*(1)*_E_*(1) */|f_B_|*^2^, equals the synchrony of the *∈*^(1)^ time series themselves, *ρ_E_*(1)*_E_*(1), times the timescale-specific strength of influence of *∈*^(1)^ on the populations, *|f_P_*(1) |^2^, and modulated by the autoregressive nature of population dynamics, 1*/|f_B_|*^2^. The term for the direct Moran effects of *∈*^(2)^ is interpreted similarly.

The components of the interacting Moran effects term in our theory are also interpretable. The phases of the quantities *f_P_*(1) and *f_P_*(2) quantify the lags in the population influences of *∈*^(1)^ and *∈*^(2)^, represented on Fig. 2 by *l_e_*_1_ and *l_e_*_2_, relative to the timescale *σ* = 1*/f* . The lag *l_n_* on

Fig. 2 manifests in the theory through the phase of *ρ_E_*(1)*_E_*(2) . The environmental effect alignment measure, *l_e_*_1_ *− l_e_*_2_ + *l_n_*, corresponds to the phase of the expression 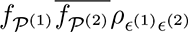, because the phase of this product equals the phase of *f_P_*(1) (which corresponds to *l_e_*_1_), minus the phase of *f_P_*(2) (which corresponds to *l_e_*_2_), plus the phase of *ρ_E_*(1)*_E_*(2) (which corresponds to *l_n_*). The real part of 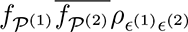 is positive whenever the phase of this quantity is close to zero, e.g., when environmental influences are positive and *l_e_*_1_ *− l_e_*_2_ + *l_n_* is close to 0 (Fig. 2a, b); and is increasingly negative as the phase of 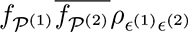 gets close to *π*, e.g., when *l_e_*_1_ *− l_e_*_2_ + *l_n_* is close to *−σ/*2 (Fig. 2c, d). The factor 1*/|f_B_|*^2^ again captures how the intrinsic nature of population dynamics modulates Moran influences.

## 3 Methods

### 3.1 Theoretical case studies

We now describe how we pursue goal G2 of the Introduction, to develop three theoretical case studies that illuminate the intuition of interacting Moran effects. For all cases, the model time step was assumed to be one quarter (q), i.e., four time steps per year. This makes no real mathematical difference, but was done to facilitate later comparisons with results for kelp data, which were sampled quarterly. For case study A (henceforth CaseA), the environmental variable *∈*^(1)^ is assumed to have a simple positive effect on populations, but lagged by one model time step (1q), i.e., *p*^(1)^ *>* 0, and *p*^(1)^ = 0 for *i /*= 1. For case study B (henceforth CaseB), *∈*^(1)^ is again assumed to have a simple positive effect on populations, but now lagged by 3q, so that *p*^(1)^ *>* 0 while *p*^(1)^ = 0 for *i /*= 3. For both CaseA and CaseB, the effects of *∈*^(2)^ are assumed to be un-lagged and positive, i.e., *p*^(2)^ *>* 0 and *p*^(2)^ = 0 for *i >* 0. For case study C (CaseC), the effects of *∈*^(1)^ are positive and lagged by 1q and the effects of *∈*^(2)^ are unlagged and negative, i.e., *p*^(1)^ *>* 0 while *p*^(1)^ = 0 for *i /*= 1, and *p*^(2)^ *<* 0 while *p*^(2)^ = 0 for *i /*= 0. The noise process (*∈*^(1)^, *∈*^(2)^) is assumed to be a Gaussian white-noise process for both CaseA and CaseB. So the random variables (*∈*^(1)^(*t*), *∈*^(2)^(*t*)) are independent and identically distributed (iid) for distinct times, *t*. Noise was positively correlated across space, and the components of *∈*^(1)^(*t*) were positively correlated with those of *∈*_*i*_^(2)^(*t*). For CaseC, the noise processes *∈*^(1)^ and *∈*^(2)^ are each assumed to exhibit spatially synchronous periodic oscillations of period one year, i.e., four model time steps, but with different phases. Peaks in *∈*^(1)^ are assumed to either lead or lag peaks in *∈*^(2)^ by 1q (we consider two sub-cases, CaseC1 and CaseC2). Such a situation could be realized by annually periodic environmental fluctuations sampled quarterly, e.g., wave action in central CA peaks annually in the winter, whereas surface-water nitrate concentrations also fluctuate with period one year, but peak in the spring, a delay of 1q compared to the wave peak. Details of the noise processes are in SI section S3. For all case studies, the autoregressive order of population dynamics is assumed, for simplicity, to be 1, i.e, *b*_1_ */*= 0 but *b_i_* = 0 for *i >* 1. Case studies do not cover the full range of possible scenarios which can be illuminated by our general theory; they were selected for the intuition they provide, and for the correspondence of some of the cases to the kelp examples we later discuss.

### 3.2 Kelp examples

To help illustrate our theory, for goal G3 of the Introduction, we also apply the theory to an exceptional dataset on giant kelp (*Macrocystis pyrifera*) dynamics off the CA coast, and associated environmental measurements. The data are based on a subset of those used by Castorani *et al*. (2022) and Walter *et al*. (2022), and data details are given in those papers and in SI section S6; Castorani *et al*. (2022) and Walter *et al*. (2022) also provide an introduction to kelp ecology, and information on why giant kelp is an excellent species for studies of synchrony. We here summarize the format of the data after a preparation and cleaning process was implemented. After preparation, kelp data consisted of 224 quarterly kelp abundance time series from locations along the CA coast, each time series spanning from quarter 1 of 1987 to quarter 4 of 2019. Time series were grouped into three regions which were analyzed separately: a more northerly central CA group of 82 locations (called CCA1); a more southerly central CA group of 82 locations (CCA2); and a group of 60 locations from southern CA, close to Santa Barbara (called SB; see Fig. 4). Each kelp measurement is an estimate of mean quarterly kelp canopy biomass (kg wet) per unit useable habitat (m^2^) along a 500m stretch of coastline. We used coastline segments where kelp was persistent through essentially all of the 1987-2019 period (SI section S6). We also had estimates of maximum wave height and mean nitrate concentration for each quarter and location. Both waves and nutrients influence kelp dynamics and synchrony (Cavanaugh *et al*., 2013; Castorani *et al*., 2022).

**Figure 4:**
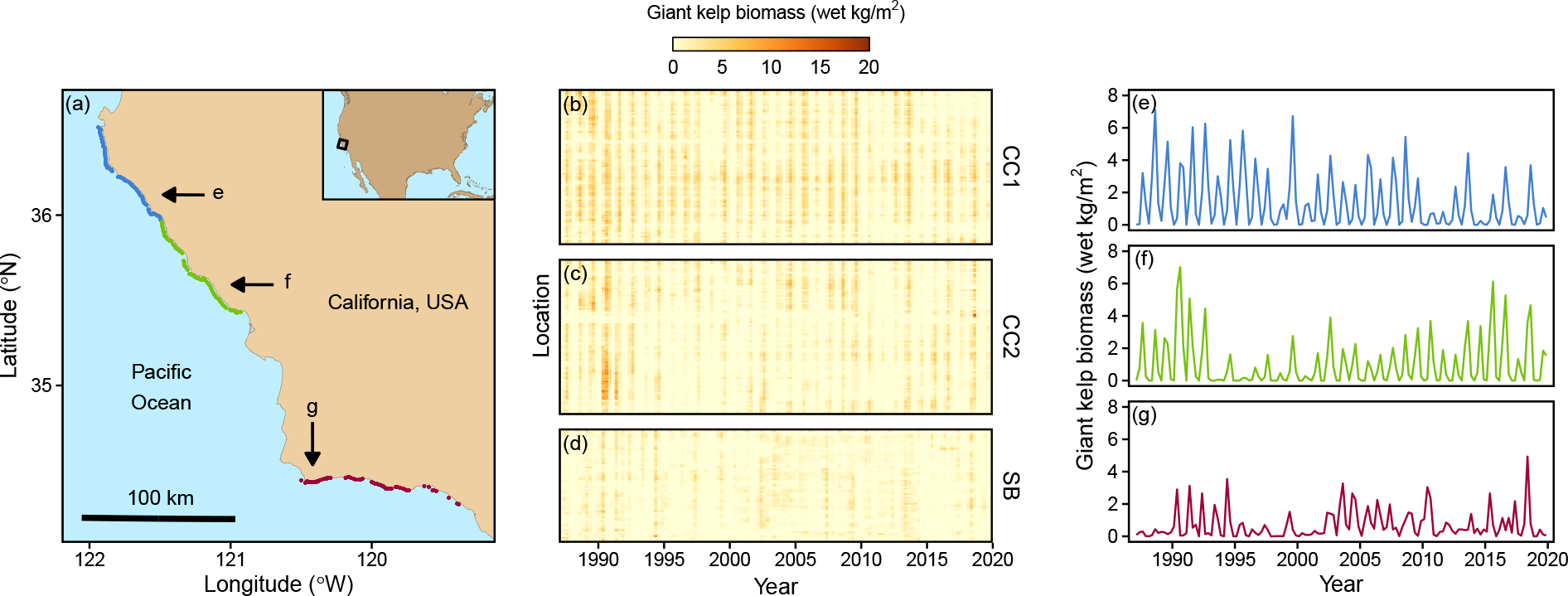
Kelp sampling sites and example time series. Sampling sites (a) were from three regions, central California 1 (CCA1, blue points), central california 2 (CCA2, green points) and the region around Santa Barbara (SB, red points). Kelp density in 500m coastline segments is shown with color intensity in (b)-(d), and those panels correspond to the regions. One example time series from each region is shown in (e)-(g), with locations at which these time series were measured labeled on panels (a)-(d). See the online version for color renderings of all figures.

Coefficients of the model (1)-(4) were separately selected for each of our three regions using linear regression methods and building on extensive prior work on the drivers of kelp dynamics. A no-intercept regression model of kelp at time *t* in location *i*, i.e., *w_i_*(*t*) in (1), against lagged values of kelp (*w_i_*(*t − l*) for *l* = 1*, … , n* in (1)), lagged and unlagged values of nitrates (*∈*^(1)^(*t − l*) for *l* = 0*, … , m*_1_ in (2)), and values of waves (*∈*^(2)^(*t − l*) for *l* = 0*, … , m*_2_ in (3)) was fitted using standard regression methods. The same regression coefficients were used for all locations within a region. Here, the *w_i_*represent linearly detrended kelp time series in one of our regions, *∈*^(1)^ were detrended nitrate time series in the region, and *∈*^(2)^ were detrended wave disturbance time series. Waves influence kelp dynamics through direct disturbance events which can damage kelp or extirpate kelp locally when waves are large (Cavanaugh *et al*., 2011; Bell *et al*., 2015; Schiel & Foster, 2015). Thus wave effects are immediate and *m*_2_ = 0 was used. Nitrates are known to fuel rapid kelp growth, though in some areas effects appear delayed by about 1q because our kelp data quantify canopy (surface) biomass, and it can take time for subsurface kelp to grow back to the surface (Cavanaugh *et al*., 2011; Bell *et al*., 2015; Schiel & Foster, 2015). Therefore *m*_1_ = 1 was used. Kelp holdfasts on the sea floor can last for multiple years, so kelp lag effects were considered: we used *n* = 4, 8, 12q in separate analyses, with this choice making no substantive difference to results (see Results). Fitted regression coefficients determined the quantities *b*_1_*, … , b_n_*, *p*^(1)^, *p*^(1)^, and *p*^(2)^ in (1)-(3), and therefore *f_B_*, *f*_P_ (1) and *f* in (5), for each of our three regions.To estimate the components *ρ_ww_*, *ρ_E_*(1)*_E_*(1) , *ρ_E_*(2)*_E_*(2) and *ρ_E_*(1)*_E_*(2) in (5), we applied the definitions of these quantities (General Theory), which required estimation from data of the spectra and cross spectra *S_wiwj_*and *S_E_*(*a*)*_E_*(*b*) for *i, j* = 1*, … , N* and *a, b* = 1, 2, where *N* is the number of locations in the region being considered. Spectral quantities were computed using the consistent estimator of section 7.4 of Brillinger (2001). The estimator is a smoothed periodogram, with the width of the smoothing kernel selected to increase with the square root of time series length. Theory was interpreted in relation to kelp ecology and theoretical case studies by plotting the components of (5) for each of our regions.

Data for the project are publicly archived (Bell *et al*., 2021). All computations were done in R version 3.6.3 on a laptop running Ubuntu Linux 16.04. Complete codes for the project workflow are at https://github.com/reumandc/InteractingMoranEffects.git.

## 4 Results

### 4.1 Illustrating properties of Moran interactions: case studies

To begin fulfilling goal G2 of the Introduction, our theoretical case studies demonstrate that interaction effects between Moran drivers: 1) can be comparable in strength to direct Moran effects; 2) can be either synergistic or destructive; and 3) can depend strongly on timescale. First, for all of our case studies, the magnitude of interaction effects was comparable to that of direct Moran effects (Fig. 5, compare the dashed and solid lines). Thus interactions can contribute substantially to overall synchrony. Second, in contrast to direct Moran effects, which are positive, interactions can be negative or positive. CaseA and CaseB showed negative interaction effects on short timescales (Fig. 5a,c); CaseC1 showed negative interactions on long timescales (Fig. 5e); and CaseC2 showed negative interactions on all timescales (Fig. 5g). Thus interaction effects can either augment or reduce synchrony. Finally, interaction effects depended strongly on timescale for all case studies. This result complements earlier studies that showed direct Moran effects can depend on timescale (Defriez *et al*., 2016; Sheppard *et al*., 2016; Desharnais *et al*., 2018; Anderson *et al*., 2021). The results of this paragraph also follow from Sheppard *et al*. (2019) and Castorani *et al*. (2022), though not straightforwardly, and those papers are case studies, whereas our results provide general theory.

**Figure 5:**
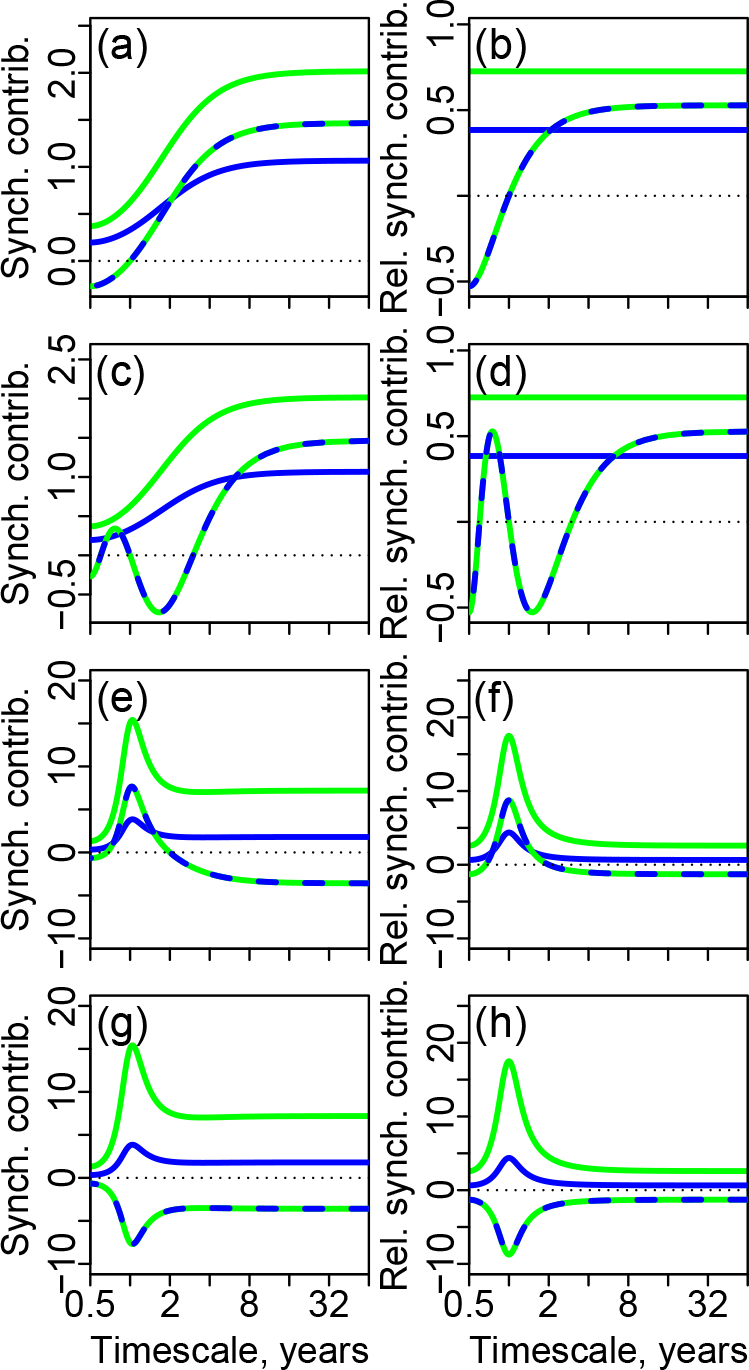
Theoretical case studies A (panels a-b), B (c-d), C1 (e-f) and C2 (g-h). Left panels (a, c, e, g) show the terms on the right side of (5). On those panels, the green line is 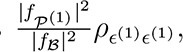, quantifying population synchrony due to the direct Moran effects of *c*^(1)^. The blue line is 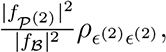, quantifying population synchrony due to the direct Moran effects of *c*^(2)^. The dashed line is 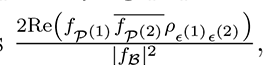, quantifying population synchrony due to interacting Moran effects. The functions plotted on b, d, f, h are those on a, c, e, g, respectively, times *|f_B_|*^2^, plotted to illustrate how autoregressive population effects modulate synchrony [see (5)]. Model parameters were: *N* = 5 and *b*_1_ = 0.4 for all case studies; = 1.1 and *p*^(2)^ = 0.8 for case study A; *p*^(1)^ = 1.1 and *p*^(2)^ = 0.8 for case study B; *p*^(1)^ = 3, *p*^(2)^ = *−*1.5 for case study C. For C1, peaks in the periodic noise process *c*^(2)^ lagged peaks in the periodic process *c*^(1)^ by 1 quarter (e-f), and for C2, *c*^(1)^ lagged *c*^(2)^ by the same amount (g-h). See SI section S3 and Fig. S1 for parameters associated with the noise for each case study, and see Methods and SI section S7 for additional details. Synch. contrib. = Synchrony contribution refers to contributions to synchrony of the individual terms in our theory; Rel. synch. contrib. = Relative synchrony contribution refers to contributions expressed without accounting for the influence of autoregressive population effects. See the online version for color renderings of all figures.

### 4.2 Building intuition for Moran interactions: case studies

CaseA helps provide an intuitive understanding about how lags in Moran drivers can produce contrasting interactions between Moran effects on different timescales, and how our theory captures that contrast. For CaseA, *∈*^(1)^ effects on populations were lagged by 1q but *∈*^(2)^ effects were unlagged (Methods). So, in the language of Fig. 2, *l_e_*_1_ = 1q and *l_e_*_2_ = 0q. The between-noise lag, *l_n_*, of Fig. 2 is 0q because (*∈*^(1)^, *∈*^(2)^) was taken to be a white noise process (see SI section S3 for details). Thus, the environmental effect alignment measure, *l_e_*_1_ *− l_e_*_2_ + *l_n_*, equals 1q. What determines the sign of interactions between Moran effects for this example is how this measure compares to the timescale/period, *σ*, being considered. On the shortest timescales (*σ* = 0.5y = 2q; Fig. 5a, left side of panel), 1q is half the period, so interaction effects are negative. On long timescales (e.g., *σ >* 8y), 1q is a negligible portion of the period, so *l_e_*_1_ *− l_e_*_2_ + *l_n_* is close to zero, relative to that period, and interaction effects are positive: relative to long timescales, *∈*^(1)^ and *∈*^(2)^ effects happen close to simultaneously, so the two noise variables reinforce each other. Comparing Fig. 5a and b shows the additional influence of autoregressive population effects, which simply multiply all Moran influences by the same timescale-dependent non-negative quantity, not altering their relative importance or sign [see (5)].

CaseB reveals how Moran effects can interact when lags are longer than 1 model time step. For CaseB, recall that *∈*^(1)^ effects on populations were lagged by 3q but *∈*^(2)^ effects were unlagged (Methods), so *l_e_*_1_ = 3q and *l_e_*_2_ = 0q in the language of Fig. 2. As for CaseA, because (*∈*^(1)^, *∈*^(2)^) was a white noise process for CaseB, *l_n_* = 0, so the environmental effect alignment measure is 3q. Interaction effects were again negative on short timescales (*σ* = 0.5y = 2q; Fig. 5c, far left side of panel) because *l_e_*_1_ *− l_e_*_2_ + *l_n_* = 3q was 1.5*σ* on that timescale, and so effects of *∈*^(1)^ and *∈*^(2)^ were in a half-phase relationship and counteracted each other. Similar to CaseA, *l_e_*_1_ *− l_e_*_2_ + *l_n_* = 3q was again negligible compared to long timescales, so interaction effects were positive on long timescales (Fig. 5c, right side of panel), though timescales had to be a bit longer than in CaseA for this approximation to be as good (compare the rates at which the dashed lines level off in Fig. 5a, c). The quantity *l_e_*_1_ *− l_e_*_2_ + *l_n_* = 3q exactly equaled the timescale for *σ* = 3q = 0.75y and equaled half the timescale for *σ* = 6q = 1.5y, hence interactions effect were, respectively, positive and negative for these timescales (Fig. 5c).

CaseA and CaseB assumed white noise processes, and therefore *l_n_* was 0. But CaseC illustrates what can happen when noise processes have lagged associations, e.g., both processes oscillate with annual periodicity and distinct phenology, an important scenario because seasonality is common.

Recall that, for CaseC, *∈*^(1)^ effects on populations were positive and lagged by 1q, i.e., *l_e_*_1_ = 1 in the language of Fig. 3; whereas *∈*^(2)^ effects were unlagged and negative, i.e., *l_e_*_2_ = 0 (Methods). We compare CaseC to the Fig. 3 *β* scenarios instead of the Fig. 2 *α* scenarios because *∈*^(2)^ effects were negative for CaseC, the situation considered by the *β* scenarios. For CaseC1, peaks in the periodic noise process *∈*^(1)^ were set up to lag peaks in the periodic process *∈*^(2)^ by 1q (Methods, SI section S3), so that *l_n_* = 1. Thus the environmental effect alignment measure, *l_e_*_1_ *− l_e_*_2_ + *l_n_*, equals 2q = 0.5y, and, on annual timescales, lags compounded, similar to Fig. 3c, d: the annual positive population effects of *∈*^(1)^ were exactly 2q offset from the annual negative effects of *∈*^(2)^, one quarter because of the lag of *∈*^(1)^ peaks behind *∈*^(2)^ peaks, and one additional quarter because of the delayed influence of *∈*^(1)^ peaks on populations. This produced reinforcing interactions between the Moran effects of *∈*^(1)^ and *∈*^(2)^ on annual timescales, as reflected by our theory (Fig. 5e,f). Contrastingly, for CaseC2, peaks in the periodic noise process *∈*^(2)^ were set up to lag peaks in the periodic process *∈*^(1)^ by 1q, so that *l_n_* = *−*1. Thus *l_e_*_1_ *− l_e_*_2_ + *l_n_* = 0 and, on annual timescales, lags cancelled, similar to Fig. 3a, b: the annual positive population effects of *∈*^(1)^ coincided with the negative effects of *∈*^(2)^ every year, because *∈*^(1)^ peaks came 1q ahead of *∈*^(2)^ peaks each year, but population influences of *∈*^(1)^ were delayed by 1q. This produced negative interactions between the Moran influences of *∈*^(1)^ and *∈*^(2)^ on annual timescales, as also reflected by our theory (Fig. 5g,h). On long timescales, interactions were the same for C1 and C2, and were negative, because, on those timescales, subannual lags make negligible difference, and the effects of *∈*^(1)^ on populations were positive and those of *∈*^(2)^ were negative.

### 4.3 Illustrating properties of Moran interactions: kelp examples

We now apply our theory to kelp, fulfilling goal G3 of the Introduction. Kelp results confirm the earlier theoretical results, based on our case studies (text above and Fig. 5), that interaction effects between Moran drivers: 1) can be comparable in strength to direct Moran effects; 2) can be either synergistic or destructive; and 3) can depend strongly on timescale. First, for all of our regions and for essentially all timescales, the magnitude of interaction effects was comparable to that of direct Moran effects (Fig. 6 for the CCA1 and SB regions and Fig. S2 for the CCA2 region). Second, interaction effects could be positive (e.g., annual timescales, CCA1 region on Fig. 6a), or negative (e.g., annual timescales, SB region on Fig. 6d; or long timescales *>* 8y for either region, Fig. 6c,f). Finally, interactions depended on timescale, e.g., for CCA1 they were positive on annual timescales (Fig. 6a) and negative on timescales *>* 8y (Fig. 6c).

**Figure 6:**
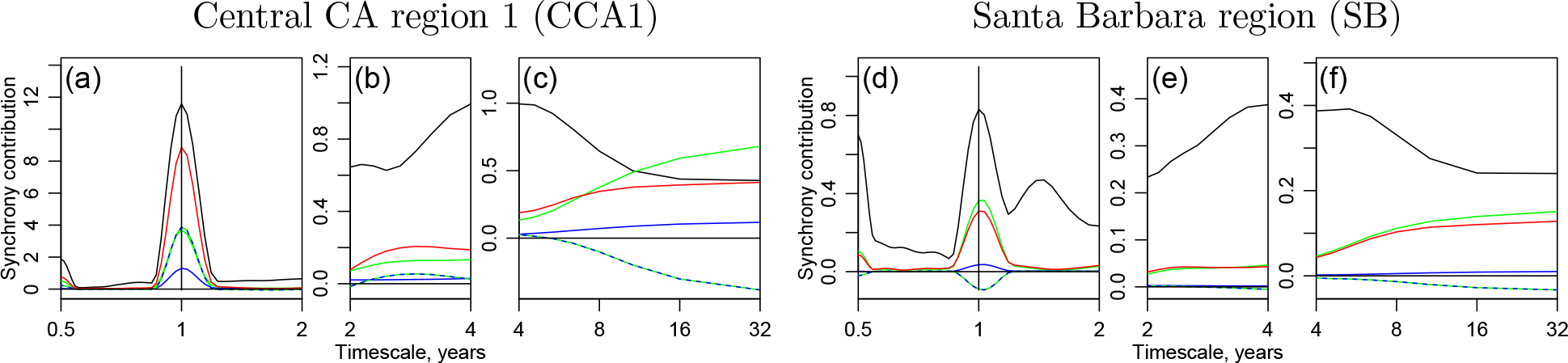
The new theory as applied to kelp, central CA 1 (CCA1) region (a-c) and Santa Barbara (SB) region (d-f). Panels show the terms on the right side of (5). Green lines show 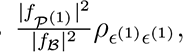, quantifying kelp population synchrony due to the direct Moran effects of *c*^(1)^, which, in this context, is nitrates. The blue line is 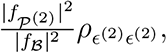, quantifying kelp population synchrony due to the direct Moran effects of *c*^(2)^, which, in this context, is waves. The dashed green-blue line is 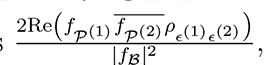, which is population synchrony due to interacting Moran effects. The red line is the sum of the green, blue and green-blue lines, and is the portion of synchrony explained by nitrates, waves, and their interactions. Explained synchrony does not equal total kelp synchrony (black line) because other, unmeasured factors also help synchronize kelp dynamics. The timescale bands 0.5 *−* 2, 2 *−* 4 and *>* 4 are separated on different panels because of the very different *y*-axis ranges. The CCA1 results approximately parallel the results for theoretical case study C scenario 1 (Fig. 5e; see text for details). See Fig. S2 for the central CA 2 (CCA2) region, for which results were substantially the same as for CCA1. This figure used kelp lag *n* = 4 [see (1)]; see Figs S3 and S4 for *n* = 8, 12, respectively. See the online version for color renderings of all figures.

### 4.4 Direct Moran effects in the kelp examples

Direct Moran effects for kelp and how they manifest in our theory are fairly straightforward. Nitrates are henceforth identified with *∈*^(1)^ and waves with *∈*^(2)^. Both nitrates and waves fluctuate seasonally in CA (Schiel & Foster, 2015). Thus nitrate and wave synchrony had a strong annual component (Fig. 7a,b,j,k for the CCA1 and SB regions, Fig. S5 for CCA2), which produced some of the annual synchrony observed in kelp (Fig. 6a,d, green and blue lines). Nitrates and waves are also synchronous on long timescales *>* 8y (Fig. 7g,h,p,q for the CCA1 and SB regions, Fig. S5 for CCA2), possibly due to the North Pacific Gyre Oscillation (Castorani *et al*., 2022; DiLorenzo *et al*., 2008). The long-timescale synchrony in nitrates and waves produced some of the long-timescale synchrony in kelp (Fig. 6c,f, green and blue lines). Kelp synchrony was stronger in CCA1 than in SB (Fig. 6a-c v. d-f, black lines) in part because the synchrony of nitrates, and of waves, was more pronounced in CCA1 than in SB (Fig. 7), and also because waves had a stronger influence on kelp in central California than in SB: regression coefficients determining *f_P_*(1), *f_P_*(2) and *f_B_* are in Table S1.

**Figure 7:**
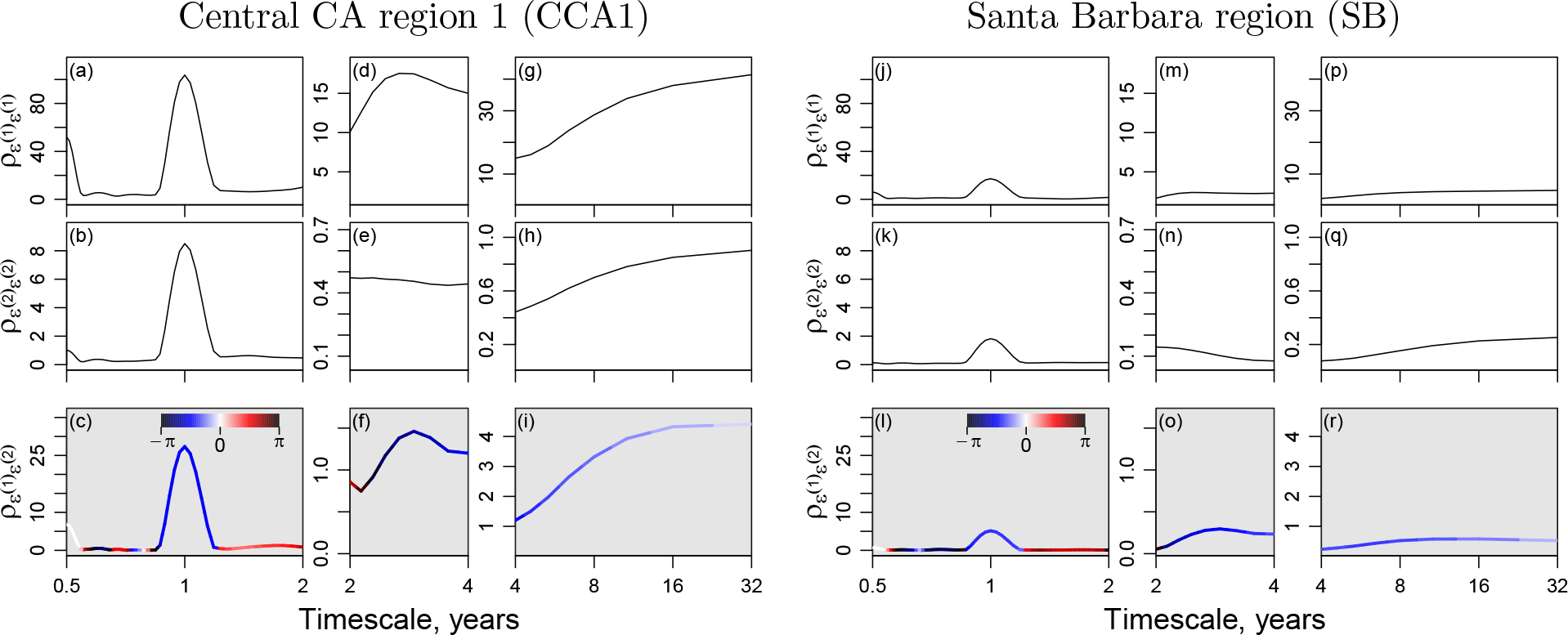
Synchrony (top two rows of panels) and cross synchrony (bottom panels) between environmental variables influencing kelp for CCA1 (a-i) and SB (j-r). Nitrates are identified with *c*^(1)^ and waves with *c*^(2)^. Vertical axis extents are the same for corresponding panels for the two regions, to facilitate comparisons. Cross synchrony is complex valued, with magnitude plotted on the vertical axis and phase displayed using color. See Figs S5 and S8 for CCA2, for which results were similar to CCA1. This figure used kelp lag *n* = 4; see Figs S6, S9, S7 and S10 for *n* = 8 and 12. See the online version for color renderings of all figures.

### 4.5 Building intuition for Moran interactions: kelp examples

Interactions between Moran effects in CCA1 were parallel to one of the theoretical case studies, CaseC1. In CCA1, nitrates had 1q-delayed positive effects on observed kelp populations and waves had immediate negative effects (Cavanaugh *et al*., 2011; Bell *et al*., 2015; Schiel & Foster, 2015), just as *∈*^(1)^ effects in CaseC1 were positive and delayed by 1q and *∈*^(2)^ effects were negative and immediate. The delayed effects of nitrates in CCA1 are reflected in Table S1, where regression results for *p*^(1)^ are close to 0 and those for *p*^(1)^ are positive. Immediate negative effects of waves in CCA1 are also reflected in Table S1, where entries for *p*^(2)^ are negative. Nitrate effects on kelp are probably delayed by a quarter in central CA because winter waves commonly remove kelp, and it takes time for kelp to grow to the surface and become detectable by satellite, only then revealing the effects of elevated nitrates (Cavanaugh *et al*., 2011; Bell *et al*., 2015; Schiel & Foster, 2015). In CCA1, annual peak nitrates tend to come in spring, whereas annual peak waves tend to come in winter (Schiel & Foster, 2015; Bell *et al*., 2015). Thus annual nitrate peaks tend to lag annual wave peaks by 1q, just as *∈*^(1)^ peaks lagged *∈*^(2)^ peaks by 1q in CaseC1. For CCA1, this is reflected in Fig. 7c, which shows that *ρ_E_*(1)*_E_*(2) has a phase of about *−π/*2 at the annual timescale. Thus *l_e_*_1_ = 1q, *l_e_*_2_ = 0q, *l_n_* = 1q, and so the environmental effect alignment measure is *l_e_*_1_ *− l_e_*_2_ + *l_n_* = 2q. Lags compound on the annual timescale, in the CCA1 case as in CaseC1: the annual positive effects of nitrates on kelp tend to come in summer, and the annual negative effects of waves come in winter, producing reinforcing interactions between Moran effects. Interactions between Moran effects on long-timescales (*>* 8y) were also similar for both CCA1 and CaseC1 (compare Figs 5e and 6c), and for the same reason in both cases: on long timescales, sub-annual lags are inconsequential, and the positive effects of nitrates/*∈*^(1)^ and the negative effects of waves/*∈*^(2)^ therefore lead to negative interactions. Interactions between Moran effects in CCA2 operated similarly to CCA1 (Fig. S8).

Due to different nitrate effects on observable kelp growth in SB compared to CCA1, interactions between Moran effects followed a different mechanism in SB compared to CCA1, leading to the slightly negative versus positive interactions already documented on annual timescales for the two regions (Fig. 6d v. a). Whereas in CCA1, nitrate effects on observed kelp density were delayed by 1q, in SB nitrate effects were observable within the same quarter: see Table S1, where *p*^(1)^ terms were close to zero for CCA1 and positive for SB, whereas *p*^(1)^ terms were positive for CCA1 and close to zero for SB. Wave effects were negative and immediate in both regions, i.e., *p*^(2)^ *<* 0 in Table S1. In southern CA, kelp are less likely to be completely removed by winter waves (Reed *et al*., 2011). Effects of elevated nitrates on the growth of kelp stands already reaching the surface can be observed within the quarter (Cavanaugh *et al*., 2011; Bell *et al*., 2015; Schiel & Foster, 2015). Annual peaks in nitrates still tended to lag annual peaks in waves by 1q in SB, as in CCA1, though the seasonal periodicity of both variables was reduced in SB compared to CCA1 (Fig. 7l v. c). Thus, in SB, the environment effect alignment measure was 1q, contrasting with its value of 2q in CCA1. Whereas in CCA1, the tendency of nitrate annual peaks to follow wave annual peaks by 1q, together with the tendency of nitrate effects on kelp to be delayed by 1q, resulted in summer positive nitrate effects and winter negative wave effects which reinforced synchrony; in SB nitrate instead both peaked and had its positive effects on kelp in spring, while waves still had negative effects on kelp in winter. Thus the effects of nitrates and waves in SB were approximately a quarter-cycle separated from each other with respect to the annual timescale, and so produced neither much reinforcement nor much destructive interference of synchrony on annual timescales (Fig. 6d). Slightly negative interactions were observed (Fig. 6d) because of slight deviations from the approximate phase alignments described above. On long timescales, interactions between Moran effects were analogous in CCA1 and SB (though weaker in SB; Fig. 6c v. f) because, again, sub-annual lags are inconsequential on long timescales and what mattered instead was the oppositely signed influences of the two environmental variables.

## 5 Discussion

We provided a general mathematical theory of the new mechanism of interactions between Moran effects. When two related spatially synchronous environmental drivers both influence a set of populations across a landscape, interactions can make synchrony in the populations either substantially stronger or markedly weaker than would otherwise be expected. Our general theory illuminates precisely how timings of influences of the drivers on populations can interact with relationships between the drivers to alter Moran effects. Interactions may vary by timescale in both their strength and sign. We used our theory and several case studies based on models and kelp populations to provide intuition about the new mechanism. Because Moran effects are ubiquitous and interactions between Moran effects were detected on both of the two occassions they have been tested for (Sheppard *et al*., 2019; Castorani *et al*., 2022), interactions may be common (also see below). Moran interactions are therefore a newly recognized and potentially widespread aspect of a fundamental means (Moran effects) by which environmental factors influence populations and diverse, synchrony-related phenomena such as ecosystem stability (Wilcox *et al*., 2017), population cycling (Anderson *et al*., 2021), and extinction (Ghosh *et al*., 2020c). Climate change is altering many aspects of environmental variables, including their means, variances and spatial correlations (Lyon *et al*., 2019; Keelings & Moradkhani, 2020), as well as relationships between environmental variables and the nature of their influences on populations. There is therefore also potential for climate change to alter interactions between Moran effects, in ways our new theory may help researchers to understand. The augmented fundamental understanding of Moran effects which we have provided may substantially benefit both basic (Liebhold *et al*., 2004) and applied (Hansen *et al*., 2020; Larsen *et al*., 2021; Herfindal *et al*., 2022) ecological research.

We argue that it is likely that interactions between Moran effect are common. Most species are influenced by more than one environmental driver. Drivers are frequently spatially autocorrelated, and are also often related to each other because of their common origin in underlying climatic phenomena such as the North Pacific Gyre Oscillation (NPGO) or El Niño Southern Oscillation (ENSO). Driver pairs which may commonly produce interacting Moran effects include instances where the same quantity is measured in distinct parts of the year (e.g., spring and summer temperatures, or March and April rainfall); or when distinct variables are measured in the same part of the year (e.g., spring temperature and precipitation). Such scenarios involving seasonality of effects, which were recently explored by Walter *et al*. (In review), can produce specific manifestations of the general mechanisms explored here. Walter *et al*. (In review) found interactions and cross synchrony to be important. Future work should systematically investigate cross synchrony (*ρ_E_*(1)*_E_*(2)) of environmental variables. Temperature and precipitation variables measured in the same season may be particularly important candidates for interactions because of the well known joint influence of these variables on plants.

We again revisit the intuition behind interacting Moran effects using the central CA kelp example as a vehicle. Large winter waves have immediate negative effects on kelp in central CA, whereas the positive effects of spring nitrates manifest in summer. So nutrient and wave effects can reinforce each other in producing annual oscillations: large kelp increases in summer due to abundant nutrients can be followed by big crashes in winter due to waves, both factors combining to accentuate the annual cycle. Thus positive interactions between Moran effects on annual timescales occur whenever years with above-average waves coincide with years with plentiful nutrients in other locations: if a large-wave year in location A coincides with a high-nutrient year in location B, both locations will tend to have bigger annual fluctuations that year, accentuating annual-timescale synchrony between the locations (Castorani *et al*., 2022). Sub-annual lags and delays make essentially no difference, however, on long timescales. On long timescales, large-wave and abundant-nutrient years counteract each other: if a multi-year period of large waves in location A coincides with a multi-year period of abundant nutrients in location B, the multi-year-average kelp abundance in A will tend to be reduced, whereas the multi-year-average kelp abundance in B will tend to be augmented, reducing long-timescale synchrony. Lags and interactions between drivers must always be compared to the timescale of interest to determine interaction effects, as in Figs 2 and 3, and as captured formally in our theory. Thus interactions between Moran effects provide yet another reason, among many reasons previous work has already documented (e.g., Vasseur & Gaedke, 2007;

Keitt, 2008; Vasseur *et al*., 2014; Sheppard *et al*., 2016; Defriez *et al*., 2016; Defriez & Reuman, 2017a,b; Desharnais *et al*., 2018; Walter *et al*., 2017; Anderson *et al*., 2021; Zhao *et al*., 2020), that patterns of synchrony must be considered from a timescale-specific viewpoint for full understanding.

Our results about kelp were consistent with those of Castorani *et al*. (2022), though that study uses distinct methods; our results complement the results of Castorani *et al*. (2022) in important ways. In spite of numerous methodological choices which differed between the two studies, both our results and Castorani *et al*. (2022) show positive effects of both nutrients and waves on synchrony on both annual and long (*>* 4y) timescales in both central and southern CA. And both studies show positive interactions between nutrient and wave Moran effects on annual timescales in central CA, but negative interactions in southern CA on annual timescales and in both central and southern CA on long timescales. The Fourier approach of our study was designed to facilitate mathematical examination of interactions between Moran effect as a general mechanism, and development of general theory; whereas the wavelet approach of Castorani *et al*. (2022) was instead optimized for detecting interactions and identifying Moran mechanisms in data, in spite of non-stationarity and other complicating features which are present in many ecological datasets. The study of Sheppard *et al*. (2019) developed the wavelet methods applied by Castorani *et al*. (2022); an open-source implementation of these methods (Reuman *et al*., 2021) can facilitate future work. The modelling approach of this study relates indirectly to the approach of Anderson *et al*. (2021), though that study concerned different research questions.

It has been a frequent topic of research why populations are often less synchronous, or sychronous over a smaller spatial extent, than might be expected given the strength and extent of synchrony of an environmental driver (Herfindal *et al*., 2022). Our new mechanism of interacting Moran effects provides both a new means by which populations may be less synchronous than population drivers; and also a new means by which populations can be more synchronous than environmental drivers. Previously known mechanisms by which populations can be less synchronous than environmental drivers include demographic stochasticity and measurement error. Antagonistic interactions between Moran drivers may be a common and previously unrecognized additional mechanism contributing to this discrepancy. On the other hand, two recent papers descibed “enhanced Moran effects” by which specific patterns of temporal autocorrelation in Moran drivers can theoretically cause greater synchrony in populations than in drivers (Massie *et al*., 2015; Desharnais *et al*., 2018). Synergistic interaction between Moran drivers are another mechanism by which populations can be more synchronous than expected.

Climate change has the potential to influence interactions between Moran effects in two specific ways which can be illuminated by our theory, and this potential should be investigated in future work. Examining the third term of (5), climate change could alter interaction effects if it: 1) alters the term *ρ_E_*(1)*_E_*(2) quantifying cross synchrony between Moran drivers; or 2) alters one of the terms *f_P_*(1) or *f_P_*(2) specifying the nature of the influence of the environmental variables *∈*^(1)^ and *∈*^(2)^, respectively, on populations. As advocated above, the term *ρ_E_*(1)*_E_*(2) should be systematically computed in future work, for a variety of environmental variables, to assess whether interactions between Moran effects are likely to be general. As part of that process, the potential for changes in *ρ_E_*(1)*_E_*(2) could also be assessed, by using either time-windowed versions or wavelet adaptations of this statistic, applied to either long-term climate records or future climate scenarios. Differences in *f_P_*(1) were responsible for differences in the nature of Moran interactions between central CA and southern CA (Results), specifically differences between the two regions in the lag of nitrate effects on kelp populations. If climate change modifies environmental effects on populations in a related way it would be expected to produce similarly large changes in Moran interactions. Climate change may alter lags and delays associated with environmental effects on populations in at least two ways: 1) by altering species phenology; and 2) by increasing or decreasing growth rates and thereby decreasing or increasing delays. Though it is too early to conclude that effects on Moran interactions are among the most important impacts of climate change, we feel the mechanisms outlined above are sufficient to warrant further investigation.

We have focussed on interactions between two Moran drivers, but synchrony in most systems may be a phenomenon with multiple (more than two) interacting causes. Kendall *et al*. (2000) considered interactions between dispersal and a Moran mechanism of synchrony. Their research questions were therefore distinct from ours, but combining their viewpoints and ours may lead to future work about interactions between dispersal and more than one Moran driver of synchrony. Although we considered only two Moran drivers in our theory and examples, essentially all population systems are influenced by multiple environmental drivers, and environmental drivers very commonly are associated with large scale climatic phenomena such as ENSO, and hence are associated with each other. Thus it may be quite common for synchrony to simultaneously be caused by dispersal and multiple distinct Moran effects, and these influences may interact in multifarious combinations. It may be necessary in future work to consider interactions between dispersal and two Moran drivers. It may be useful to consider cases for which multiple related Moran drivers all interact. Dispersal can readily be added to our modelling framework: Desharnais *et al*. (2018) performed a spectral analysis on a model similar to ours which included dispersal. Future work should consider whether and when scenarios of multi-driver interactions between causes of synchrony can lead to synchrony patterns which differ fundamentally from what one would expect from one or two mechanisms. Potential interactions increase as the square of the number of drivers, so interactions seem likely to become more important as our viewpoint of synchrony expands.

## Supporting information

Complete supporting information

## Acknowledgments

This study was partly funded by the U.S. National Science Foundation (NSF) through linked NSFOCE awards 2023555, 2023523, 2140335, and 2023474 to M.C.N.C, K.C.C., T.W.B, and D.C.R., respectively; and by NSF-Math Bio award 1714195 to D.C.R.; and by support to D.C.R. from the James S. McDonnell Foundation, the Humboldt Foundation, and the California Department of Fish and Wildlife Delta Science Program. This project used data developed through the Santa Barbara Coastal Long Term Ecological Research project, funded through NSF-OCE award 1831937. The authors thank Vadim Karatayev, Maowei Liang, Kyle Emery, Nat Coombs, Adeola Adeboje and Ethan Kadiyala for helpful discussions.

