## Supplementary material for "How environmental drivers of spatial synchrony interact": Complete supporting information

### Contents

**S1 Mathematical details of the example in the Introduction** **2**

**S2 Measuring synchrony** **2**

**S3 Examples of cross synchrony** **3**

**S4 Model specification, full details** **5**

**S5 Derivation of the main equation** **6**

**S6 Details of data and data cleaning** **8**

**S7 Details of the theoretical case studies** **8**

### List of Tables

S1 Regression coefficients used to parameterize (1)-(4) of the main text for each of our three regions. All values rounded to two decimal places. . . . . . . . . . . . . . . . . . . . . . . . 18

### S1 Mathematical details of the example in the Introduction

In the case of two environmental variables, the Moran theorem implies that population synchrony is

$$\begin{aligned}\mathrm{cor}(w_1, w_2) &= \mathrm{cor}(\epsilon_1^{(1)} + \epsilon_1^{(2)}, \epsilon_2^{(1)} + \epsilon_2^{(2)}) && (1)\\ &= \frac{\mathrm{cov}(\epsilon_1^{(1)}, \epsilon_2^{(1)}) + \mathrm{cov}(\epsilon_1^{(1)}, \epsilon_2^{(2)}) + \mathrm{cov}(\epsilon_1^{(2)}, \epsilon_2^{(1)}) + \mathrm{cov}(\epsilon_1^{(2)}, \epsilon_2^{(2)})}{\sqrt{2 + 2\mathrm{cov}(\epsilon_1^{(1)}, \epsilon_1^{(2)})}\sqrt{2 + 2\mathrm{cov}(\epsilon_2^{(1)}, \epsilon_2^{(2)})}}, && (2)\\ & && (3)\end{aligned}$$

which depends not only on the standard environmental synchrony measures $\mathrm{cov}(\epsilon_1^{(a)}, \epsilon_2^{(a)})$, for $a = 1, 2$, but also on the "cross synchrony" measures $\mathrm{cov}(\epsilon_1^{(a)}, \epsilon_2^{(b)})$ for $a \neq b$.

### S2 Measuring synchrony

If $w = (w_1, \ldots, w_N)$ is an $N$-dimensional stationary stochastic process, where the index $i = 1, \ldots, N$ corresponds to sampling locations, then we measure the synchrony of $w$ as

$$\rho_{ww} = \left(\frac{1}{N^2 - N}\right) \sum_{i \neq j} S_{w_i w_j}, \tag{4}$$

where $S_{ww}$ is the spectral matrix of the process $w$. This is a function of frequency or timescale because $S_{w_i w_j}$ is a function of frequency or timescale. To see that $\rho_{ww}$ is a real-valued function, note that $\overline{S_{ww}} = S_{ww}^{\tau}$, where the superscript $\tau$ denotes matrix transpose [Brillinger (2001), theorem 2.5.1 on p. 24]. Therefore

$$\begin{aligned}\overline{\rho_{ww}} &= \left(\frac{1}{N^2 - N}\right) \sum_{i \neq j} \overline{S_{w_i w_j}} && (5)\\ &= \left(\frac{1}{N^2 - N}\right) \sum_{i \neq j} S_{w_j w_i} && (6)\\ &= \left(\frac{1}{N^2 - N}\right) \sum_{i \neq j} S_{w_i w_j} && (7)\\ &= \rho_{ww}. && (8)\end{aligned}$$

The quantity $\rho_{ww}$ makes sense as a measure of synchrony in part because

$$\begin{aligned}\rho_{ww} &= \mathrm{Re}(\rho_{ww}) && (9)\\ &= \left(\frac{1}{N^2 - N}\right) \sum_{i \neq j} \mathrm{Re}(S_{w_i w_j}). && (10)\end{aligned}$$

The quantity $\mathrm{Re}(S_{w_i w_j})$ is the cospectrum of the component processes $w_i$ and $w_j$, and an appropriate integral across frequencies of the cospectrum equals the covariance, $\mathrm{cov}(w_i, w_j)$ (Brillinger, 2001). Thus an integral of $\rho_{ww}$ across frequencies equals

$$\left(\frac{1}{N^2 - N}\right) \sum_{i \neq j} \mathrm{cov}(w_i, w_j), \tag{11}$$

which is a standard quantity that can be used to measure non-frequency-specific synchrony. Thus $\rho_{ww}$ is a frequency-specific generalization of a classic, non-frequency-specific measure of synchrony.

If $\epsilon^{(1)} = (\epsilon_1^{(1)}, \ldots, \epsilon_N^{(1)})$ and $\epsilon^{(2)} = (\epsilon_1^{(2)}, \ldots, \epsilon_N^{(2)})$ are two $N$-dimensional stationary stochastic processes, we define the *cross-variable spatial synchrony* (or, simply, the *cross synchrony*) between $\epsilon^{(1)}$ and $\epsilon^{(2)}$ as

$$\rho_{\epsilon^{(1)}\epsilon^{(2)}} = \left(\frac{1}{N^2 - N}\right) \sum_{i \neq j} S_{\epsilon_i^{(1)} \epsilon_j^{(2)}}. \tag{12}$$

This is a complex-valued function of frequency or timescale. Why is this interpretable as cross-variable spatial synchrony? A quantity representing spatial synchrony should compare values of the processes $\epsilon^{(1)}$ and $\epsilon^{(2)}$ in one location to values in another location, since spatial synchrony is about relationships between distinct locations; hence our sum is over $i \neq j$. A quantity representing cross-variable synchrony should make comparisons between two variables, as opposed to comparisons between the same variable in different locations; hence we use the cross spectrum between variable 1 in location $i$ ($\epsilon_i^{(1)}$) and variable 2 in location $j$ ($\epsilon_j^{(2)}$). Finally, a quantity representing cross-variable synchrony needs to include the possibility of time delays in the relationship between the two variables. For instance, if $\epsilon_j^{(2)}$ is a time-lagged version of $\epsilon_i^{(1)}$, but the effects of $\epsilon_j^{(2)}$ on a biotic process $w_j$ are immediate, whereas the effects of $\epsilon_i^{(1)}$ on $w_i$ are delayed by the same lag, then the effects of $\epsilon_i^{(1)}$ on $w_i$ will be correlated with the effects of $\epsilon_j^{(2)}$ on $w_j$, i.e., this combination of properties of $\epsilon^{(1)}$ and $\epsilon^{(2)}$ will produce synchrony, in the classic sense of correlation, between the biotic processes $w_i$ and $w_j$. Thus we use the cross spectrum, $S_{\epsilon_i^{(1)} \epsilon_j^{(2)}}$, for which the phase angle at frequency $f$ measures the time delay between frequency-$f$ oscillations of $\epsilon_i^{(1)}$ and frequency-$f$ oscillations of $\epsilon_j^{(2)}$.

#### S3 Examples of cross synchrony

The following examples should help clarify the properties of our definition of cross synchrony. For pedagogical clarity, we start with a white-noise example that does not illustrate the full value of the idea of cross synchrony, but which is simple; we then move on to an example which is more complex but also more illustrative. These examples are also used in the theoretical case studies (main text).

For the simple example, to be used in theoretical case studies A and B (Methods), we assume $(\epsilon^{(1)}(t), \epsilon^{(2)}(t))$ are drawn, independently for each $t$, from a $2N$-dimensional multivariate normal distribution with mean $(0, ..., 0)$ and covariance matrix $\Sigma$, where the entries $\Sigma_{ij}$ of $\Sigma$ are as follows. For $i = j$, $\Sigma_{ij} = 1$. For $i \neq j$ and $1 \leq i, j \leq N$, $\Sigma_{ij}$ equals a parameter $v_{11}$. For $i \neq j$ and $N+1 \leq i, j \leq 2N$, $\Sigma_{ij}$ equals a second parameter $v_{22}$. The remaining entries of $\Sigma$ equal a third parameter, $v_{12}$. The parameters $v_{11}$, $v_{22}$ and $v_{12}$ must be chosen to yield a positive definite matrix $\Sigma$.

Now, having specified the noise process $(\epsilon^{(1)}, \epsilon^{(2)})$, we derive the quantities $\rho_{\epsilon^{(1)}\epsilon^{(1)}}$, $\rho_{\epsilon^{(2)}\epsilon^{(2)}}$, and $\rho_{\epsilon^{(1)}\epsilon^{(2)}}$. For $a$ and $b$ equal to 1 or 2, the formula $S_{\epsilon_i^{(a)} \epsilon_j^{(b)}}(f) = \sum_{u=-\infty}^{\infty} c_{\epsilon_i^{(a)} \epsilon_j^{(b)}}(u) \exp(-2\pi\iota f u)$, where $c_{\epsilon_i^{(a)} \epsilon_j^{(b)}}(u) = \mathrm{cov}(\epsilon_i^{(a)}(t+u), \epsilon_j^{(b)}(t))$, follows from definitions of the spectrum and cross spectrum, given, for instance, in 2.5.2 on p. 23 of Brillinger (2001). The quantity $c_{\epsilon_i^{(a)} \epsilon_j^{(b)}}(u) = \mathrm{cov}(\epsilon_i^{(a)}(t+u), \epsilon_j^{(b)}(t))$ is independent of $t$

for any stationary process $(\epsilon^{(1)}, \epsilon^{(2)})$, and in particular for the temporally iid process that we assumed to operate here. Furthermore, because $\epsilon_i^{(a)}(t+u)$ and $\epsilon_j^{(b)}(t)$ were assumed independent for $u \neq 0$, we have

$$S_{\epsilon_i^{(a)}\epsilon_j^{(b)}}(f) = c_{\epsilon_i^{(a)}\epsilon_j^{(b)}}(0) = \begin{cases} 1 & \text{if } a = b \text{ and } i = j \\ v_{11} & \text{if } a = b = 1 \text{ and } i \neq j \\ v_{22} & \text{if } a = b = 2 \text{ and } i \neq j \\ v_{12} & \text{if } a \neq b. \end{cases} \tag{13}$$

Thus the spectra and cross spectra in this example are constant. It is then easy to see that $\rho_{\epsilon^{(1)}\epsilon^{(1)}} = v_{11}$, $\rho_{\epsilon^{(2)}\epsilon^{(2)}} = v_{22}$, and $\rho_{\epsilon^{(1)}\epsilon^{(2)}} = v_{12}$, based on the definitions of these quantities given previously. For theoretical case studies A and B in the main text, we used $N = 5$, $v_{11} = v_{22} = 0.6$ and $v_{12} = 0.3$.

These results make intuitive sense, for a white-noise process, for the following reasons. First, $\rho_{\epsilon^{(1)}\epsilon^{(1)}}$ and $\rho_{\epsilon^{(2)}\epsilon^{(2)}}$, which are automatically real-valued (SI section S2), should be flat for a white noise process, i.e., independent of timescale, because spectral quantities are flat for time series with no temporal dependencies. For the same reason, $\rho_{\epsilon^{(1)}\epsilon^{(2)}}$ should be flat. It makes sense that cross synchrony in this example is real-valued because a complex-valued $\rho_{\epsilon^{(1)}\epsilon^{(2)}}$ would imply lagged or phase-delayed relationships between $\epsilon^{(1)}$ and $\epsilon^{(2)}$, not possible with a white-noise process.

For a more illustrative example, used in case study C (Methods), the noise process $(\epsilon^{(1)}, \epsilon^{(2)})$ is generated via a three-step process. First, assume $(\tilde{\zeta}^{(1)}(t), \tilde{\zeta}^{(2)}(t))$ are drawn, independently for each $t$, from a $2N$-dimensional multivariate normal distribution with mean $(0, \ldots, 0)$ and covariance matrix $\Psi$, where the entries $\Psi_{ij}$ of $\Psi$ are as follows. For $i = j$, $\Psi_{ij} = 1$. For $i \neq j$ and $1 \leq i, j \leq N$, $\Psi_{ij}$ equals a parameter $w_{11}$. For $i \neq j$ and $N + 1 \leq i, j \leq 2N$, $\Psi_{ij}$ equals a second parameter $w_{22}$. The remaining entries of $\Psi$ equal a third parameter, $w_{12}$. The parameters $w_{11}$, $w_{22}$ and $w_{12}$ must again be chosen to yield a positive definite matrix. Second, let $\zeta^{(1)}(t) = \tilde{\zeta}^{(1)}(t - l_1)$ and let $\zeta^{(2)}(t) = \tilde{\zeta}^{(2)}(t - l_2)$ for fixed integers $l_1$ and $l_2$. Finally, let $\epsilon_i^{(a)}(t) = c_1\epsilon_i^{(a)}(t-1) + c_2\epsilon_i^{(a)}(t-2) + \zeta_i^{(a)}(t)$ for $i = 1, \ldots, N$ and $a = 1, 2$. The second step above ensures that $\epsilon^{(1)}$ and $\epsilon^{(2)}$ are related to each other in a time-delayed manner. The third step ensures that $\epsilon^{(1)}$ and $\epsilon^{(2)}$ can oscillate with a dominant periodic component, depending on the values of $c_1$ and $c_2$ used: the autoregressive order-two process used can produce dynamics with a peaked spectrum, as we will see below (see also theorem 1 below).

To compute $\rho_{\epsilon^{(1)}\epsilon^{(1)}}$, $\rho_{\epsilon^{(2)}\epsilon^{(2)}}$ and $\rho_{\epsilon^{(1)}\epsilon^{(2)}}$ for this example, we first compute $S_{\epsilon^{(1)}\epsilon^{(1)}}$, $S_{\epsilon^{(2)}\epsilon^{(2)}}$ and $S_{\epsilon^{(1)}\epsilon^{(2)}}$ using theorem 1 below. Letting $\epsilon = (\epsilon^{(1)}, \epsilon^{(2)})$ and $\zeta(t) = (\zeta^{(1)}, \zeta^{(2)})$, we have $\epsilon_i(t) = c_1\epsilon_i(t-1) + c_2\epsilon_i(t-2) + \zeta_i(t)$ for $i = 1, \ldots, 2N$. We then apply theorem 1 to get that $S_{\epsilon\epsilon} = \left|\frac{1}{1-c_1\mu-c_2\mu^2}\right|^2 S_{\zeta\zeta}$. Considering upper-left $N \times N$ sub-matrices gives $S_{\epsilon^{(1)}\epsilon^{(1)}} = \left|\frac{1}{1-c_1\mu-c_2\mu^2}\right|^2 S_{\zeta^{(1)}\zeta^{(1)}}$. Considering lower-right $N \times N$ sub-matrices gives $S_{\epsilon^{(2)}\epsilon^{(2)}} = \left|\frac{1}{1-c_1\mu-c_2\mu^2}\right|^2 S_{\zeta^{(2)}\zeta^{(2)}}$. Considering upper-right $N \times N$ sub-matrices gives $S_{\epsilon^{(1)}\epsilon^{(2)}} = \left|\frac{1}{1-c_1\mu-c_2\mu^2}\right|^2 S_{\zeta^{(1)}\zeta^{(2)}}$. Because the $\zeta^{(1)}(t)$ are iid for distinct times $t$, we can again use the definitions of the spectrum and the cross spectrum and reasoning similar to that used above for the previous examples to show that $S_{\zeta^{(1)}\zeta^{(1)}}$ equals the upper-left $N \times N$ sub-matrix of $\Psi$, which is a matrix with 1 in every diagonal entry and $w_{11}$ in all off-diagonal entries. Likewise, $S_{\zeta^{(2)}\zeta^{(2)}}$ equals the lower-right $N \times N$ sub-matrix of $\Psi$, which has 1 in all diagonal entries and $w_{22}$ in all off-diagonal entries. To get $S_{\zeta^{(1)}\zeta^{(2)}}$, we write $S_{\zeta_i^{(1)}\zeta_j^{(2)}}(f) = \sum_{u=-\infty}^{\infty} c_{\zeta_i^{(1)}\zeta_j^{(2)}}(u) \exp(-2\pi\iota f u)$. Here,

$$c_{\zeta_i^{(1)}\zeta_j^{(2)}}(u) = \mathrm{cov}(\zeta_i^{(1)}(t+u), \zeta_j^{(2)}(t)) \tag{14}$$

$$= \mathrm{cov}(\tilde{\zeta}_i^{(1)}(t+u-l_1), \tilde{\zeta}_j^{(2)}(t-l_2)) \tag{15}$$

$$= \begin{cases} w_{12} & \text{if } u - l_1 = -l_2 \\ 0 & \text{otherwise.} \end{cases} \tag{16}$$

So $S_{\zeta_i^{(1)}\zeta_j^{(2)}} = w_{12}\exp(-2\pi\iota f(l_1 - l_2))$. It then follows straightforwardly that $\rho_{\epsilon^{(1)}\epsilon^{(1)}} = \left|\frac{1}{1-c_1\mu-c_2\mu^2}\right|^2 w_{11}$, $\rho_{\epsilon^{(2)}\epsilon^{(2)}} = \left|\frac{1}{1-c_1\mu-c_2\mu^2}\right|^2 w_{22}$ and $\rho_{\epsilon^{(1)}\epsilon^{(2)}} = \left|\frac{1}{1-c_1\mu-c_2\mu^2}\right|^2 w_{12}\exp(-2\pi\iota f(l_1 - l_2))$. Fig. S1 displays these quantities for the two parameterizations of $(\epsilon^{(1)}, \epsilon^{(2)})$ that we use for theoretical case study C.

We now explore why the results above for $\rho_{\epsilon^{(1)}\epsilon^{(1)}}$, $\rho_{\epsilon^{(2)}\epsilon^{(2)}}$ and $\rho_{\epsilon^{(1)}\epsilon^{(2)}}$ make sense for the noise processes we described, particularly for the sets of parameters used for theoretical case study C in the main text (see Fig. S1). The pre-factor $\left|\frac{1}{1-c_1\mu-c_2\mu^2}\right|^2$ in all these expressions reflects the autoregressive manner in which the noises $\epsilon$ were defined. Autoregressive models of order 2 are well known to be capable of producing oscillations for certain values of the parameters. For instance, for case study C, we use $c_1 = 0$ and $c_2 = -0.4444$, values which cause the expression $\left|\frac{1}{1-c_1\mu-c_2\mu^2}\right|^2$ to have a single peak at frequency 0.25 cycles per sampling interval. It makes sense that the synchrony quantities $\rho_{\epsilon^{(1)}\epsilon^{(1)}}$, $\rho_{\epsilon^{(2)}\epsilon^{(2)}}$ and $\rho_{\epsilon^{(1)}\epsilon^{(2)}}$ for periodically oscillating quantities should also have a peak at the timescale of oscillation. The term $\exp(-2\pi\iota f(l_1 - l_2))$ in the expression for $\rho_{\epsilon^{(1)}\epsilon^{(2)}}$ reflects the fact that cross variable synchrony for this example is lagged by an amount $l_1 - l_2$. The lack of a similar term in the expressions for $\rho_{\epsilon^{(1)}\epsilon^{(1)}}$ or $\rho_{\epsilon^{(2)}\epsilon^{(2)}}$ reflects that fact that there are no lags built into within-variable synchrony in this example.

**Theorem 1.** *If $\zeta(t)$ is an $N$-dimensional second-order stationary ergodic stochastic process with component expected values $0$, if $\epsilon_i(t) = c_1\epsilon_i(t-1) + \cdots + c_n\epsilon_i(t-n) + q_0\zeta_i(t) + \cdots + q_m\zeta_i(t-m)$ for $i = 1, \ldots, N$, with $c_n \neq 0$ and $q_m \neq 0$, and if the complex roots of the polynomial $1 - c_1 z - c_2 z^2 - \cdots c_n z^n = 0$ have modulus greater than $1$, then $S_{\epsilon\epsilon}(f) = \left|\frac{\gamma}{1-\lambda}\right|^2 S_{\zeta\zeta}(f)$ where $\gamma = q_0 + q_1\mu + \cdots + q_m\mu^m$, $\lambda = c_1\mu + c_2\mu^2 + \cdots + c_n\mu^n$, and $\mu = \exp(-2\pi\iota f)$, $\iota$ is the imaginary unit, and $f$ is frequency in cycles per time step.*

*Proof.* This theorem is identical to theorem 2 in SI section S1.5 of Anderson *et al.* (2021), where a proof is also provided. □

### S4 Model specification, full details

The population model is specified starting with the equation

$$\begin{aligned} w_i(t) = {} & b_1 w_i(t-1) + \cdots + b_n w_i(t-n) && (17)\\ & + p_0^{(1)}\epsilon_i^{(1)}(t) + \cdots + p_{m_1}^{(1)}\epsilon_i^{(1)}(t - m_1) && (18)\\ & + p_0^{(2)}\epsilon_i^{(2)}(t) + \cdots + p_{m_2}^{(2)}\epsilon_i^{(2)}(t - m_2) && (19)\\ & + \delta_i(t), && (20) \end{aligned}$$

where $i = 1, \ldots, N$ indexes habitat patches or sampling locations, $w_i(t)$ is a measure of the population in location $i$ at time $t$, and $\epsilon^{(1)} = (\epsilon_1^{(1)}, \ldots, \epsilon_N^{(1)})$, $\epsilon^{(2)} = (\epsilon_1^{(2)}, \ldots, \epsilon_N^{(2)})$ and $\delta = (\delta_1, \ldots, \delta_N)$ are environmental processes influencing the populations. Note that this model makes some assumptions from the outset. First, it assumes a homogeneous influence of the environmental variables on the populations in the different sampling locations, i.e., the parameters $p$ are spatially homogeneous. Second, autoregressive population effects are also assumed spatially homogeneous. Third, we assume that the combined process $(\epsilon_1^{(1)}, \ldots, \epsilon_N^{(1)}, \epsilon_1^{(2)}, \ldots, \epsilon_N^{(2)}, \delta_1, \ldots, \delta_N)$ is a second-order stationary stochastic process (Brillinger, 2001) with the expected values of its component processes equal to 0. We do *not* assume that the unmeasured noise terms $(\delta_1, \ldots, \delta_N)$ are spatially asynchronous or independent through time. Finally, we assume the complex roots of the polynomial $1 - b_1 z - \cdots - b_n z^n = 0$ have modulus greater than 1, an assumption needed for the existence of a stationary stochastic process solution for the model and for invertability of the filter $\mathcal{B}$ which we will define below. See Reinsel (1997); Shumway & Stoffer (2000); Brillinger (2001) for background on the statistical theory of stationary stochastic processes.

Letting $I_N$ denote the $N \times N$ identity matrix and letting $B$ denote the backshift operator, we define the linear filters

$$\mathcal{P}^{(1)} = p_0^{(1)} I_N + p_1^{(1)} I_N B + \cdots + p_{m_1}^{(1)} I_N B^{m_1}, \tag{21}$$

$$\mathcal{P}^{(2)} = p_0^{(2)} I_N + p_1^{(2)} I_N B + \cdots + p_{m_2}^{(2)} I_N B^{m_2}, \tag{22}$$

$$\mathcal{B} = I_N - b_1 I_N B - b_2 I_N B^2 - \cdots b_n I_N B^n. \tag{23}$$

Using the language of filters and stochastic processes, the original model (17-20) can then be written as

$$\mathcal{B} w = \mathcal{P}^{(1)} \epsilon^{(1)} + \mathcal{P}^{(2)} \epsilon^{(2)} + \delta. \tag{24}$$

Writing the model in the language of stochastic processes and linear filters makes it straightforward to calculate the spectral matrix, $S_{ww}$, of the stochastic process, $w$, which we do in section S5.

### S5 Derivation of the main equation

Computing the spectral matrix of both sides of the model (24) gives

$$T(\mathcal{B}) S_{ww} T(\mathcal{B})^* = T(\mathcal{P}^{(1)}) S_{\epsilon^{(1)}\epsilon^{(1)}} T(\mathcal{P}^{(1)})^* \tag{25}$$

$$+ T(\mathcal{P}^{(2)}) S_{\epsilon^{(2)}\epsilon^{(2)}} T(\mathcal{P}^{(2)})^* \tag{26}$$

$$+ S_{\delta\delta} \tag{27}$$

$$+ T(\mathcal{P}^{(1)}) S_{\epsilon^{(1)}\epsilon^{(2)}} T(\mathcal{P}^{(2)})^* + T(\mathcal{P}^{(2)}) S_{\epsilon^{(2)}\epsilon^{(1)}} T(\mathcal{P}^{(1)})^* \tag{28}$$

$$+ T(\mathcal{P}^{(1)}) S_{\epsilon^{(1)}\delta} + S_{\delta\epsilon^{(1)}} T(\mathcal{P}^{(1)})^* \tag{29}$$

$$+ T(\mathcal{P}^{(2)}) S_{\epsilon^{(2)}\delta} + S_{\delta\epsilon^{(2)}} T(\mathcal{P}^{(2)})^*, \tag{30}$$

where $T(\mathcal{B})$, $T(\mathcal{P}^{(1)})$ and $T(\mathcal{P}^{(2)})$ are the tranfer functions of the filters $\mathcal{B}$, $\mathcal{P}^{(1)}$ and $\mathcal{P}^{(2)}$, respectively, $S_{ww}$ is the spectral matrix of the process $w$, $S_{\epsilon^{(i)}\epsilon^{(i)}}$ is the spectral matrix of the process $\epsilon^{(i)}$, $S_{\delta\delta}$ is the spectral matrix of the process $\delta$, $S_{\epsilon^{(i)}\epsilon^{(j)}}$ is the cross spectral matrix of the processes $\epsilon^{(i)}$ and $\epsilon^{(j)}$, $S_{\epsilon^{(i)}\delta}$ is the cross spectral matrix of the processes $\epsilon^{(i)}$ and $\delta$, $S_{\delta\epsilon^{(i)}}$ is the cross spectral matrix of the processes $\delta$ and $\epsilon^{(i)}$, and the superscript $*$ denotes the conjugate transpose of a matrix.

Letting $\mu = \exp(-2\pi\iota f)$ where $\iota$ is the imaginary unit and $f$ is frequency in units of cycles per time step, we have

$$T(\mathcal{B}) = (1 - b_1\mu - b_2\mu^2 - \cdots - b_n\mu^n) I_N \tag{31}$$

$$T(\mathcal{P}^{(1)}) = (p_0^{(1)} + p_1^{(1)}\mu + \cdots + p_{m_1}^{(1)}\mu^{m_1}) I_N \tag{32}$$

$$T(\mathcal{P}^{(2)}) = (p_0^{(2)} + p_1^{(2)}\mu + \cdots + p_{m_2}^{(2)}\mu^{m_2}) I_N. \tag{33}$$

These are all scalar matrices, so (25-30) simplifies to

$$S_{ww} = \frac{|f_{\mathcal{P}^{(1)}}|^2}{|f_{\mathcal{B}}|^2} S_{\epsilon^{(1)}\epsilon^{(1)}} \tag{34}$$

$$+ \frac{|f_{\mathcal{P}^{(2)}}|^2}{|f_{\mathcal{B}}|^2} S_{\epsilon^{(2)}\epsilon^{(2)}} \tag{35}$$

$$+ \frac{1}{|f_{\mathcal{B}}|^2} S_{\delta\delta} \tag{36}$$

$$+ \frac{1}{|f_{\mathcal{B}}|^2} \left[ f_{\mathcal{P}^{(1)}} \overline{f_{\mathcal{P}^{(2)}}} S_{\epsilon^{(1)}\epsilon^{(2)}} + f_{\mathcal{P}^{(2)}} \overline{f_{\mathcal{P}^{(1)}}} S_{\epsilon^{(2)}\epsilon^{(1)}} \right] \tag{37}$$

$$+ \frac{1}{|f_{\mathcal{B}}|^2} \left[ f_{\mathcal{P}^{(1)}} S_{\epsilon^{(1)}\delta} + \overline{f_{\mathcal{P}^{(1)}}} S_{\delta\epsilon^{(1)}} \right] \tag{38}$$

$$+ \frac{1}{|f_{\mathcal{B}}|^2} \left[ f_{\mathcal{P}^{(2)}} S_{\epsilon^{(2)}\delta} + \overline{f_{\mathcal{P}^{(2)}}} S_{\delta\epsilon^{(2)}} \right], \tag{39}$$

where

$$f_{\mathcal{B}} = 1 - b_1\mu - b_2\mu^2 - \cdots - b_n\mu^n \quad (40)$$

$$f_{\mathcal{P}^{(1)}} = p_0^{(1)} + p_1^{(1)}\mu + \cdots + p_{m_1}^{(1)}\mu^{m_1} \quad (41)$$

$$f_{\mathcal{P}^{(2)}} = p_0^{(2)} + p_1^{(2)}\mu + \cdots + p_{m_2}^{(2)}\mu^{m_2}. \quad (42)$$

We here made use of the assumption that the complex roots of the polynomial $1 - b_1 z - \cdots - b_n z^n = 0$ have modulus greater than 1, so that $f_{\mathcal{B}}$ is non-zero.

Now, averaging the off-diagonal entries of both sides of (34-39) gives

$$\left(\frac{1}{N^2-N}\right)\sum_{i\neq j} S_{w_i w_j} = \frac{|f_{\mathcal{P}^{(1)}}|^2}{|f_{\mathcal{B}}|^2}\left(\frac{1}{N^2-N}\right)\sum_{i\neq j} S_{\epsilon_i^{(1)}\epsilon_j^{(1)}} \quad (43)$$

$$+ \frac{|f_{\mathcal{P}^{(2)}}|^2}{|f_{\mathcal{B}}|^2}\left(\frac{1}{N^2-N}\right)\sum_{i\neq j} S_{\epsilon_i^{(2)}\epsilon_j^{(2)}} \quad (44)$$

$$+ \frac{1}{|f_{\mathcal{B}}|^2}\left(\frac{1}{N^2-N}\right)\sum_{i\neq j} S_{\delta_i\delta_j} \quad (45)$$

$$+ \frac{2}{|f_{\mathcal{B}}|^2}\left(\frac{1}{N^2-N}\right)\sum_{i\neq j} \mathrm{Re}\left(f_{\mathcal{P}^{(1)}}\overline{f_{\mathcal{P}^{(2)}}}S_{\epsilon_i^{(1)}\epsilon_j^{(2)}}\right) \quad (46)$$

$$+ \frac{2}{|f_{\mathcal{B}}|^2}\left(\frac{1}{N^2-N}\right)\sum_{i\neq j} \mathrm{Re}\left(f_{\mathcal{P}^{(1)}}S_{\epsilon_i^{(1)}\delta_j}\right) \quad (47)$$

$$+ \frac{2}{|f_{\mathcal{B}}|^2}\left(\frac{1}{N^2-N}\right)\sum_{i\neq j} \mathrm{Re}\left(f_{\mathcal{P}^{(2)}}S_{\epsilon_i^{(2)}\delta_j}\right), \quad (48)$$

where $\mathrm{Re}(\cdot)$ denotes the real part, and where (46-48) follows from two observations: that the two summands in the brackets of each of the terms (37-39) are conjugate-transpose pairs; and that, for any complex matrix, $A$, $\sum_{i\neq j}(A_{ij} + \overline{A_{ji}}) = \sum_{i\neq j}(A_{ij} + \overline{A_{ij}}) = 2\sum_{i\neq j}\mathrm{Re}(A_{ij})$. Now, making use of the definitions of section S2, this becomes

$$\rho_{ww} = \frac{|f_{\mathcal{P}^{(1)}}|^2}{|f_{\mathcal{B}}|^2}\rho_{\epsilon^{(1)}\epsilon^{(1)}} + \frac{|f_{\mathcal{P}^{(2)}}|^2}{|f_{\mathcal{B}}|^2}\rho_{\epsilon^{(2)}\epsilon^{(2)}} + \frac{1}{|f_{\mathcal{B}}|^2}\rho_{\delta\delta} \quad (49)$$

$$+ \frac{2}{|f_{\mathcal{B}}|^2}\mathrm{Re}\left(f_{\mathcal{P}^{(1)}}\overline{f_{\mathcal{P}^{(2)}}}\rho_{\epsilon^{(1)}\epsilon^{(2)}}\right) \quad (50)$$

$$+ \frac{2}{|f_{\mathcal{B}}|^2}\mathrm{Re}\left(f_{\mathcal{P}^{(1)}}\rho_{\epsilon^{(1)}\delta}\right) \quad (51)$$

$$+ \frac{2}{|f_{\mathcal{B}}|^2}\mathrm{Re}\left(f_{\mathcal{P}^{(2)}}\rho_{\epsilon^{(2)}\delta}\right), \quad (52)$$

which expresses population synchrony in terms of noise synchrony and cross synchrony. If the processes $\epsilon^{(1)}$ and $\epsilon^{(2)}$ were measured and the process $\delta$ represents the aggregate effects of unknown processes, then we can write

$$\rho_{ww} = \frac{|f_{\mathcal{P}^{(1)}}|^2}{|f_{\mathcal{B}}|^2}\rho_{\epsilon^{(1)}\epsilon^{(1)}} \quad (53)$$

$$+ \frac{|f_{\mathcal{P}^{(2)}}|^2}{|f_{\mathcal{B}}|^2}\rho_{\epsilon^{(2)}\epsilon^{(2)}} \quad (54)$$

$$+ \frac{2}{|f_{\mathcal{B}}|^2}\mathrm{Re}\left(f_{\mathcal{P}^{(1)}}\overline{f_{\mathcal{P}^{(2)}}}\rho_{\epsilon^{(1)}\epsilon^{(2)}}\right) \quad (55)$$

$$+ \text{other contributions.} \quad (56)$$

The term (53) is contributions to the synchrony of $w$ due to direct Moran effects of $\epsilon^{(1)}$. The term (54) is contributions to the synchrony of $w$ due to direct Moran effects of $\epsilon^{(2)}$. The term (55) is contributions to the synchrony of $w$ due to interactions between the Moran effects of $\epsilon^{(1)}$ and $\epsilon^{(2)}$. The "other contributions" may include direct Moran effects of other, unmeasured environmental factors, as well as interactions between these effects and Moran effects of the measured factors $\epsilon^{(1)}$ and $\epsilon^{(2)}$.

### S6 Details of data and data cleaning

Floating surface canopy of kelp was measured by remote sensing, with details and validation reported by Cavanaugh *et al.* (2011) and Bell *et al.* (2020, 2021). Multispectral Landsat imagery of 30m resolution was used. Quarterly mean biomass values and spatial extent of canopy within 500m coastline segments (henceforth called "locations") were computed by averaging. Quarters were: Q1, January-March; Q2, April-June; Q3, July-September; Q4, October-December. Data were thusly available from 437 locations spanning approximately from Monterey Bay to San Diego. We then chose locations for analysis for which kelp was "persistent", in this context meaning absent or missing data for no more than three years; this was to focus on population dynamics of kelp rather than extinction-recolonization dynamics, which occur in some locations and which can relate to sediment dynamics on the ocean floor and other factors not currently of primary interest. Of the original 437 locations, 361 (83%) were deemed persistent according to this criterion. Missing values in the remaining time series were then filled with seasonal medians. The biomass and spatial extent time series for each location were then used together to produce one time series estimating biomass density per unit useable habitat at the site, as follows: every value of the biomass time series was divided by the 90$^{\text{th}}$ quantile of the distribution of the spatial extent time series at the site. The 90$^{\text{th}}$ quantile was used instead of the maximum value of the spatial extent time series because maximum statistics have high variance. This procedure served to normalize total biomass by an estimate of total useable habitat at a site. Kelp establish on rocky substrate, and areas with sandy or sedimented substrate are not useable kelp habitat. All time series were then linearly detrended, and only those time series were utilized which were located within the three regions pictured in Fig. 4 in the main text. Regions were selected to represent distinct dynamical patterns known to occur in central and southern CA, and also to have about the same number of time series per selected region.

Wave and nutrient data were also available for each quarter at each location. Wave data were derived by applying a swell propagation model with bouy-based wave measurements and a validation techniques (Wingeart *et al.*, 2001; Hanson *et al.*, 2009; Bell *et al.*, 2015; O'Reilly *et al.*, 2016). Data on nitrates were produced (Snyder *et al.*, 2020) by exploiting an established empirical relationship between surface nitrate concentration and sea surface temperature (SST) in coastal CA (Zimmerman & Kremer, 1984; Palacios *et al.*, 2013; Snyder *et al.*, 2020) and SST data derived from Advanced Very High Resolution Radiometer satellite imagery. Wave and nutrient time series in each location were again linearly detrended before use.

### S7 Details of the theoretical case studies

The noise processes $(\epsilon^{(1)}, \epsilon^{(2)})$ used in case studies A, B and C, which were only partly specified in the main text, are fully specified in SI section S3. We next derive, for our case studies, the values of the terms that appear in (5) of the main text. For case study A, parameter specifications presented in Methods of the main text straightforwardly imply that $f_{\mathcal{B}} = 1 - b_1\mu$, $f_{\mathcal{P}^{(1)}} = p_1^{(1)}\mu$ and $f_{\mathcal{P}^{(2)}} = p_0^{(2)}$. In SI section S3, we computed that $\rho_{\epsilon^{(1)}\epsilon^{(1)}} = v_{11}$, $\rho_{\epsilon^{(2)}\epsilon^{(2)}} = v_{22}$, and $\rho_{\epsilon^{(1)}\epsilon^{(2)}} = v_{12}$.

For case study B, details presented in Methods of the main text imply that $f_{\mathcal{B}} = 1 - b_1\mu$, $f_{\mathcal{P}^{(1)}} = p_n^{(1)}\mu^n$ for $n = 3$, and $f_{\mathcal{P}^{(2)}} = p_0^{(2)}$. Because the noise process for B is taken to be the same as that for A, we again have $\rho_{\epsilon^{(1)}\epsilon^{(1)}} = v_{11}$, $\rho_{\epsilon^{(2)}\epsilon^{(2)}} = v_{22}$ and $\rho_{\epsilon^{(1)}\epsilon^{(2)}} = v_{12}$.

For case study C, again using details from Methods, $f_{\mathcal{B}} = 1 - b_1\mu$, $f_{\mathcal{P}^{(1)}} = p_{n_1}^{(1)}\mu^{n_1}$ for $n_1 = 1$, and

$f_{\mathcal{P}^{(2)}} = p^{(2)}_{n_2}\mu^{n_2}$ for $n_2 = 0$. In SI section S3, we computed that $\rho_{\epsilon^{(1)}\epsilon^{(1)}} = \left|\frac{1}{1-c_1\mu-c_2\mu^2}\right|^2 w_{11}$, $\rho_{\epsilon^{(2)}\epsilon^{(2)}} = \left|\frac{1}{1-c_1\mu-c_2\mu^2}\right|^2 w_{22}$ and $\rho_{\epsilon^{(1)}\epsilon^{(2)}} = \left|\frac{1}{1-c_1\mu-c_2\mu^2}\right|^2 w_{12}\exp(-2\pi\iota f(l_1 - l_2))$.

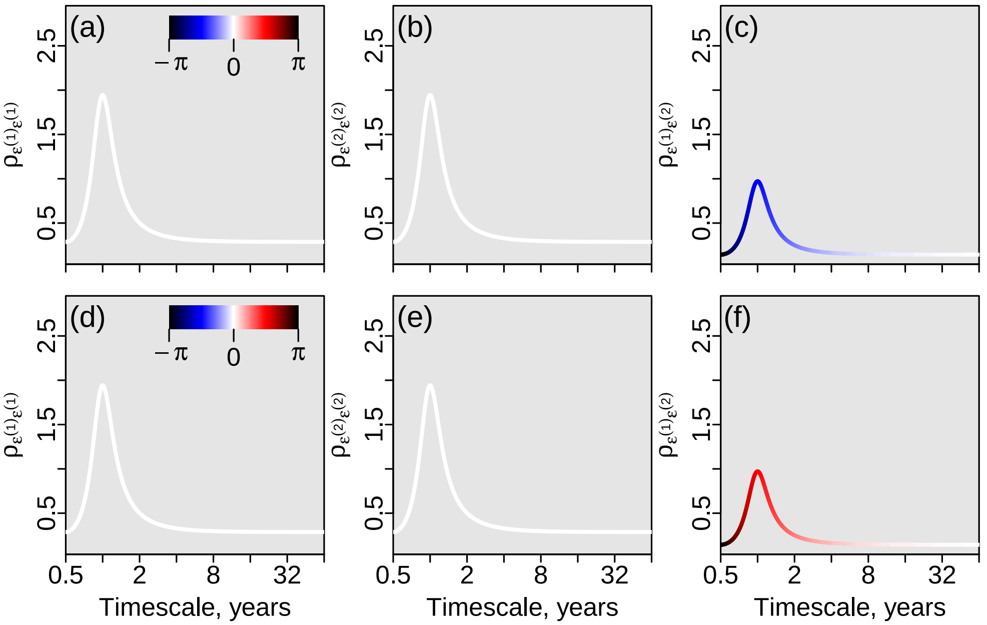

Figure S1: The quantities $\rho_{\epsilon^{(1)}\epsilon^{(1)}}$, $\rho_{\epsilon^{(2)}\epsilon^{(2)}}$, and $\rho_{\epsilon^{(1)}\epsilon^{(2)}}$ for the noise described in SI section S3 and used for theoretical case study C. These are complex-valued functions of timescale; their magnitude is displayed on vertical axes and their phase is displayed in color. Two alternative sets of parameters were used for panels a-c and d-f, respectively, and are also used in the two scenarios considered for case study C. Parameters were: $N = 5$, $c_1 = 0$, $c_2 = -0.4444$, $w_{11} = w22 = 0.6$ and $w_{12} = 0.3$ for both sets of panels; $l_1 = 1$ and $l_2 = 0$ for panels a-c; and $l_1 = 0$ and $l_2 = 1$ for panels d-f.

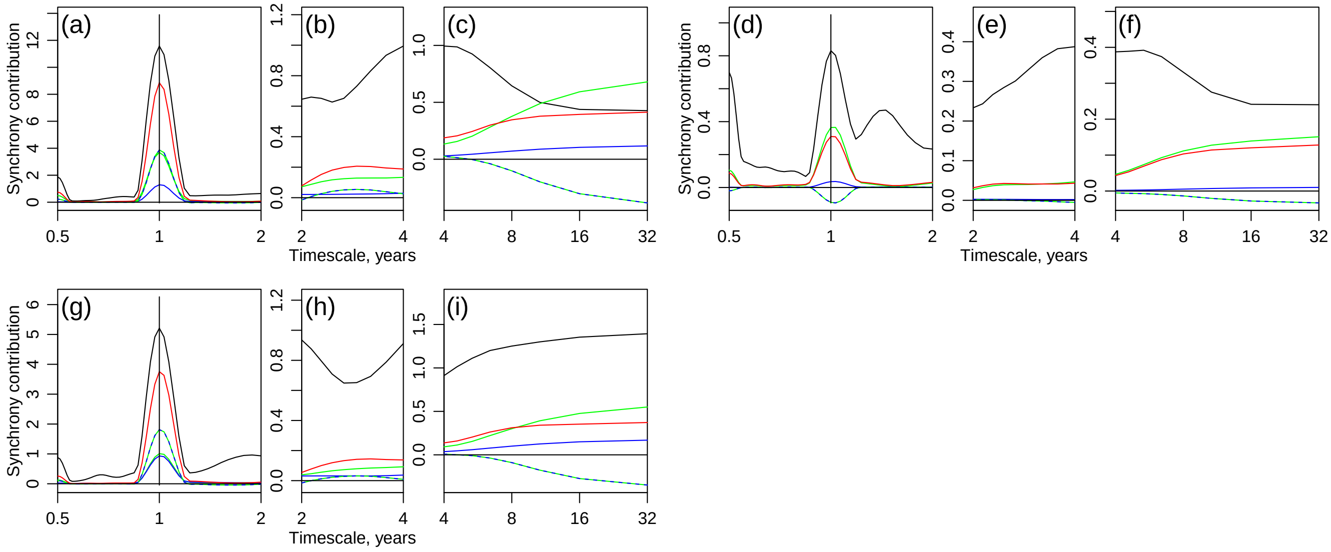

Figure S2: Same format as Fig. 6 of the main text, but including not only results for the CCA1 region (a-c) and the SB region (d-f), but also results from the CCA2 region (g-i).

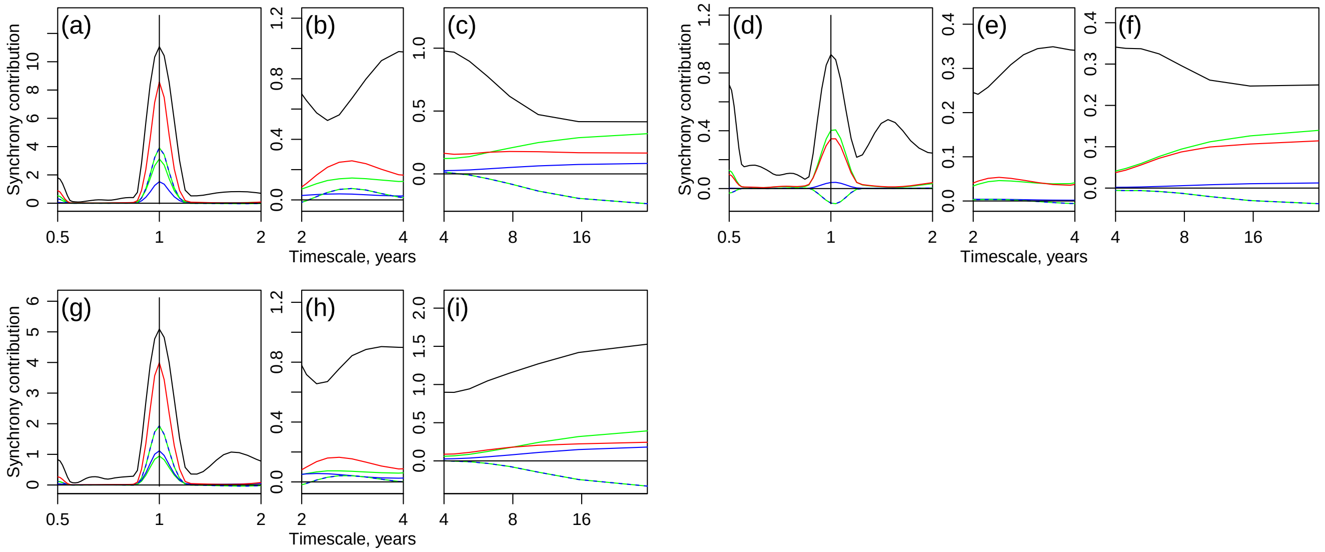

Figure S3: Same format as Fig. S2, but based on a model that considered potential kelp lag effects from up to 8 quarters in the past (see Methods). Results were substantively the same as for Fig. S2.

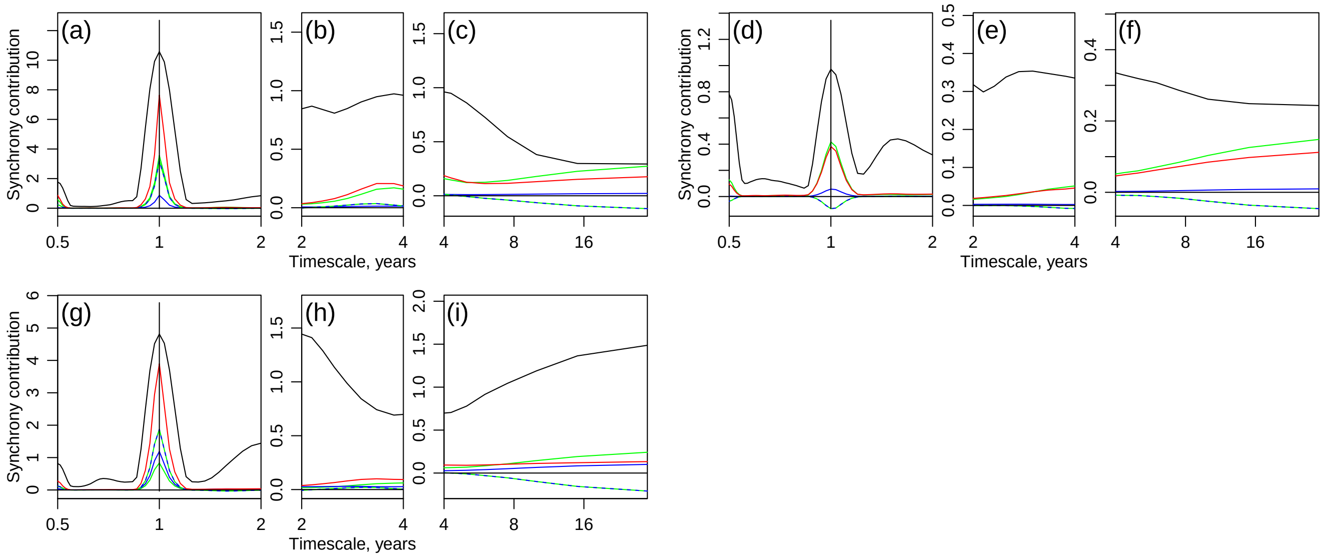

Figure S4: Same format as Fig. S2, but based on a model that considered potential kelp lag effects from up to 12 quarters in the past (see Methods). Results were substantively the same as for Fig. S2.

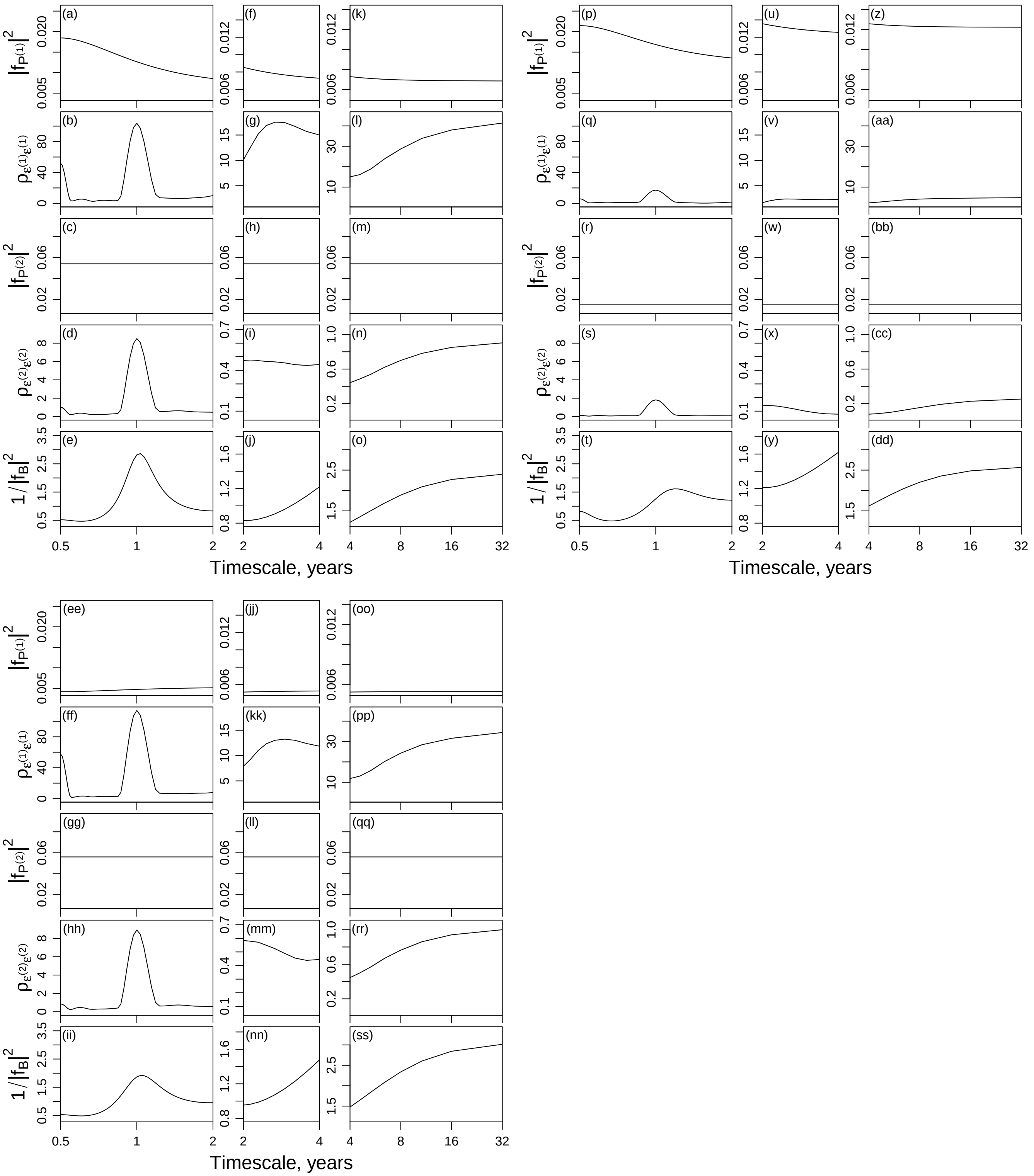

Figure S5: Plots of all constituents of the terms in theory that pertain to direct Moran effects, for kelp, for the CCA1 region (a-o), the SB region (p-dd) and the CCA2 region (ee-ss), based on a model that considered potential kelp lag effects from up to 4 quarters in the past (see Methods).

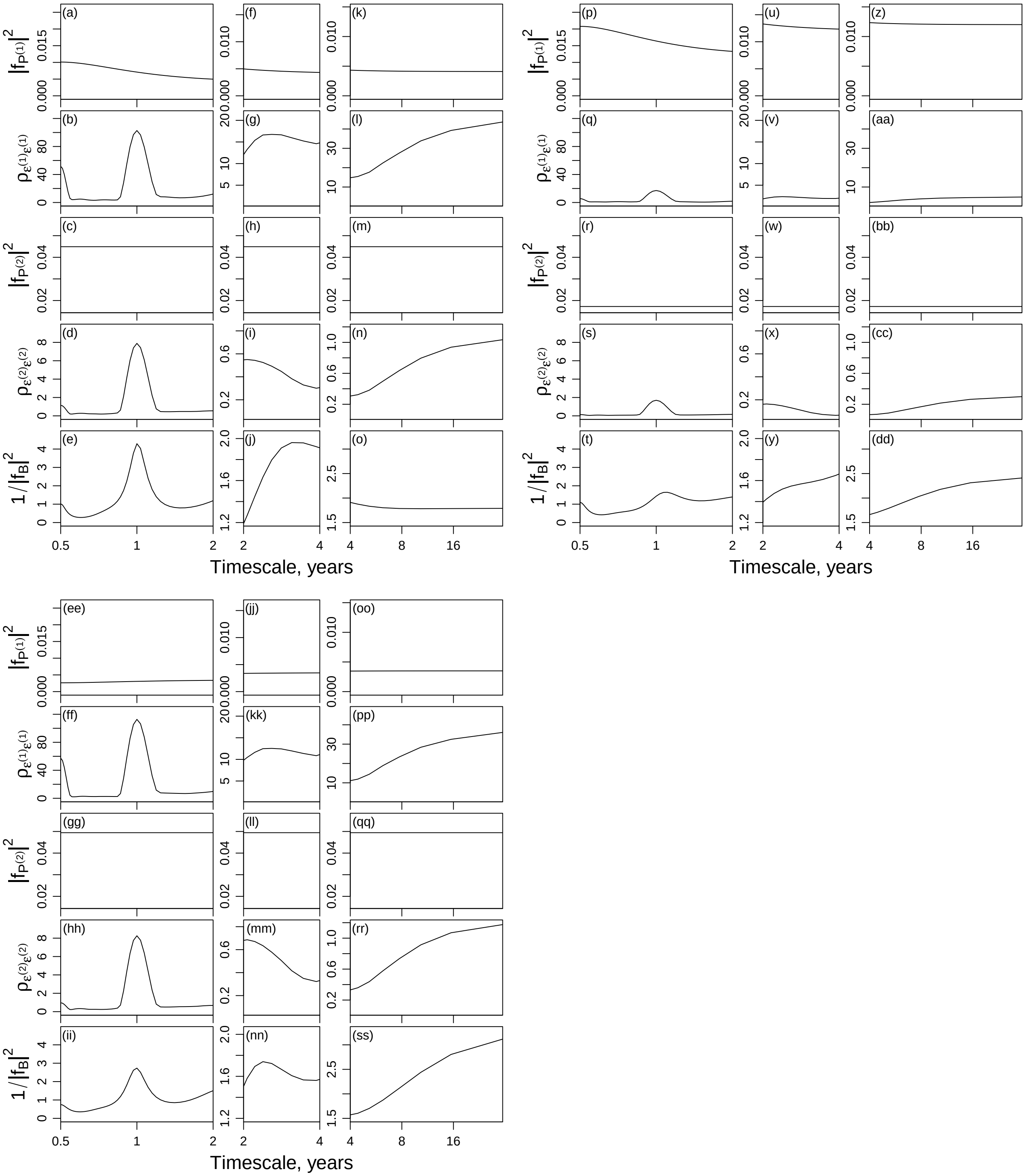

Figure S6: Same format as Fig. S5, but based on a model that considered potential kelp lag effects from up to 8 quarters in the past (see Methods). Results were substantively the same as for Fig. S5.

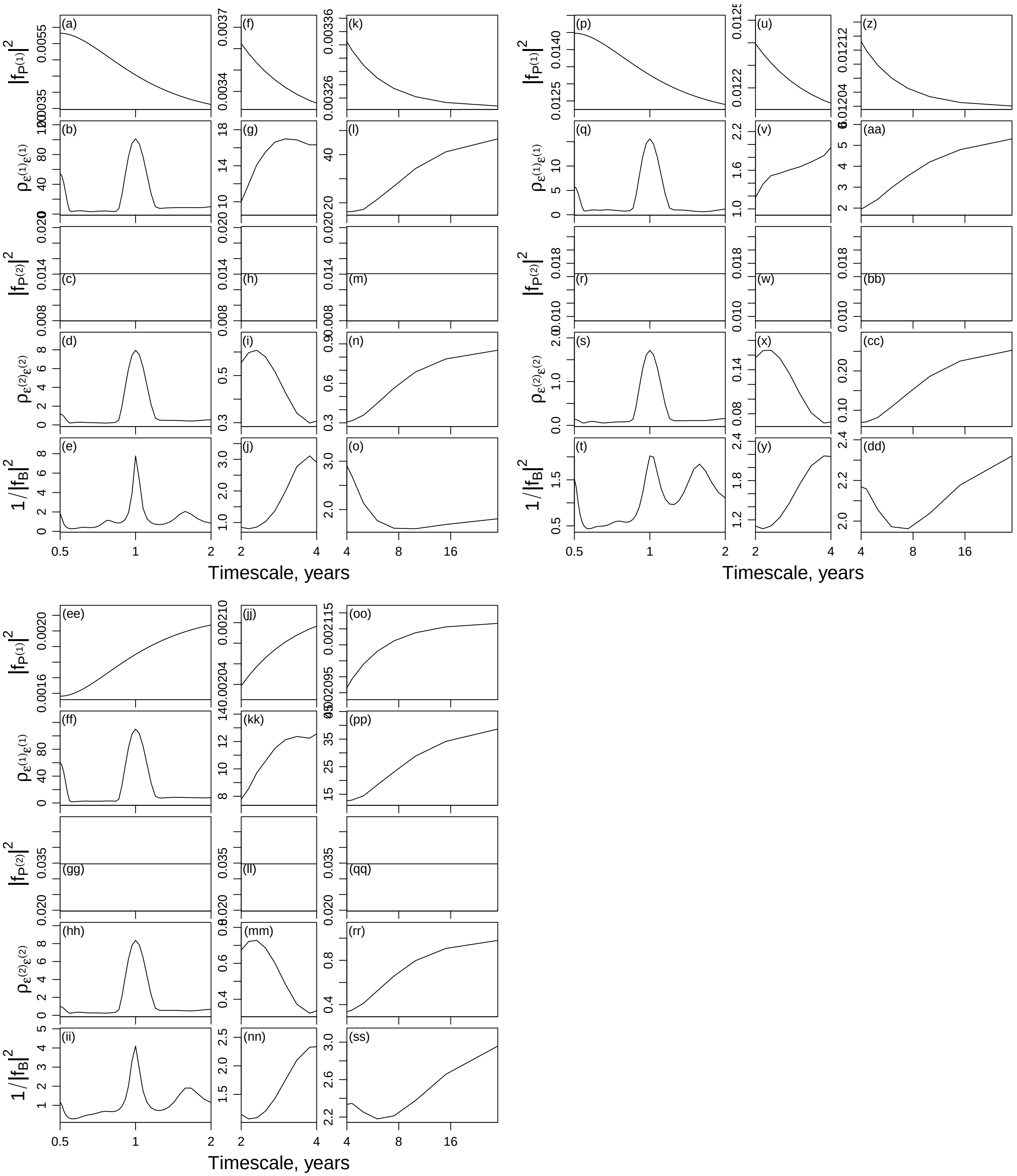

Figure S7: Same format as Fig. S5, but based on a model that considered potential kelp lag effects from up to 12 quarters in the past (see Methods). Results were substantively the same as for Fig. S5.

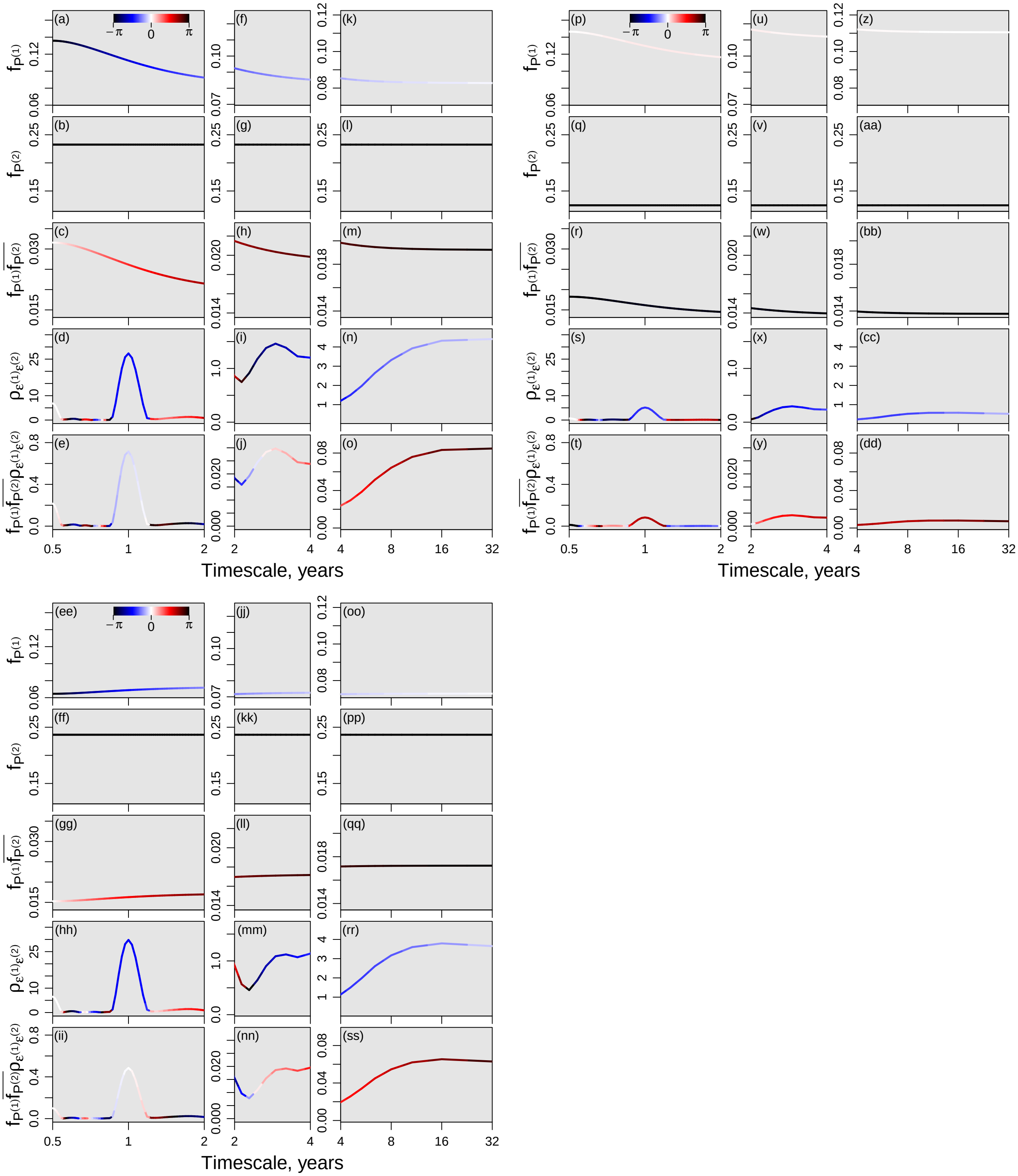

Figure S8: Plots of all constituents of the terms in theory that pertain to interactions between Moran effects, for kelp, for CCA1 region (a-o), the SB region (p-dd), and the CCA2 region (ee-ss), based on a model that considered potential kelp lag effects from up to 4 quarters in the past (see Methods).

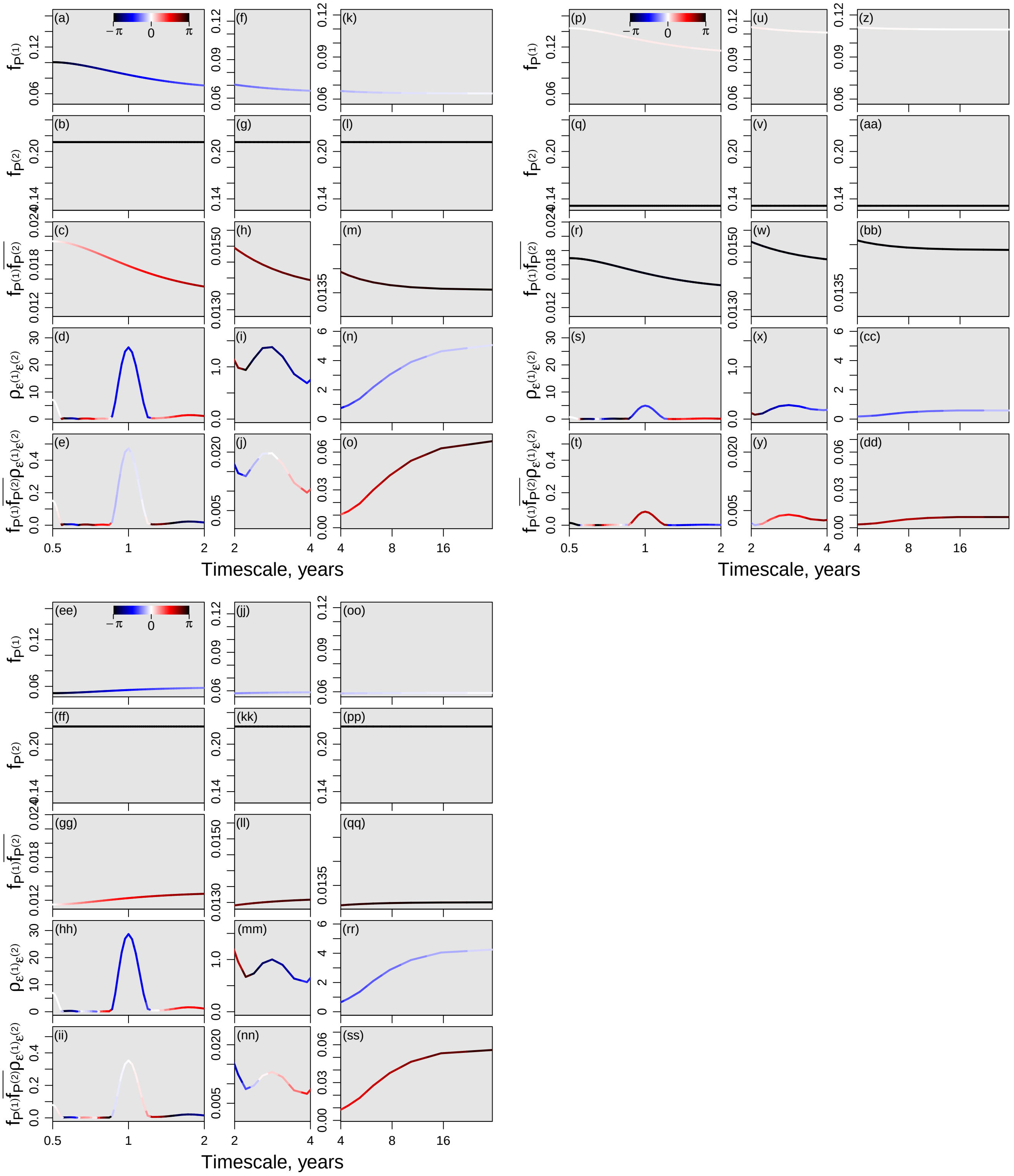

Figure S9: Same format as Fig. S8, but based on a model that considered potential kelp lag effects from up to 8 quarters in the past (see Methods). Results were substantively the same as for Fig. S8.

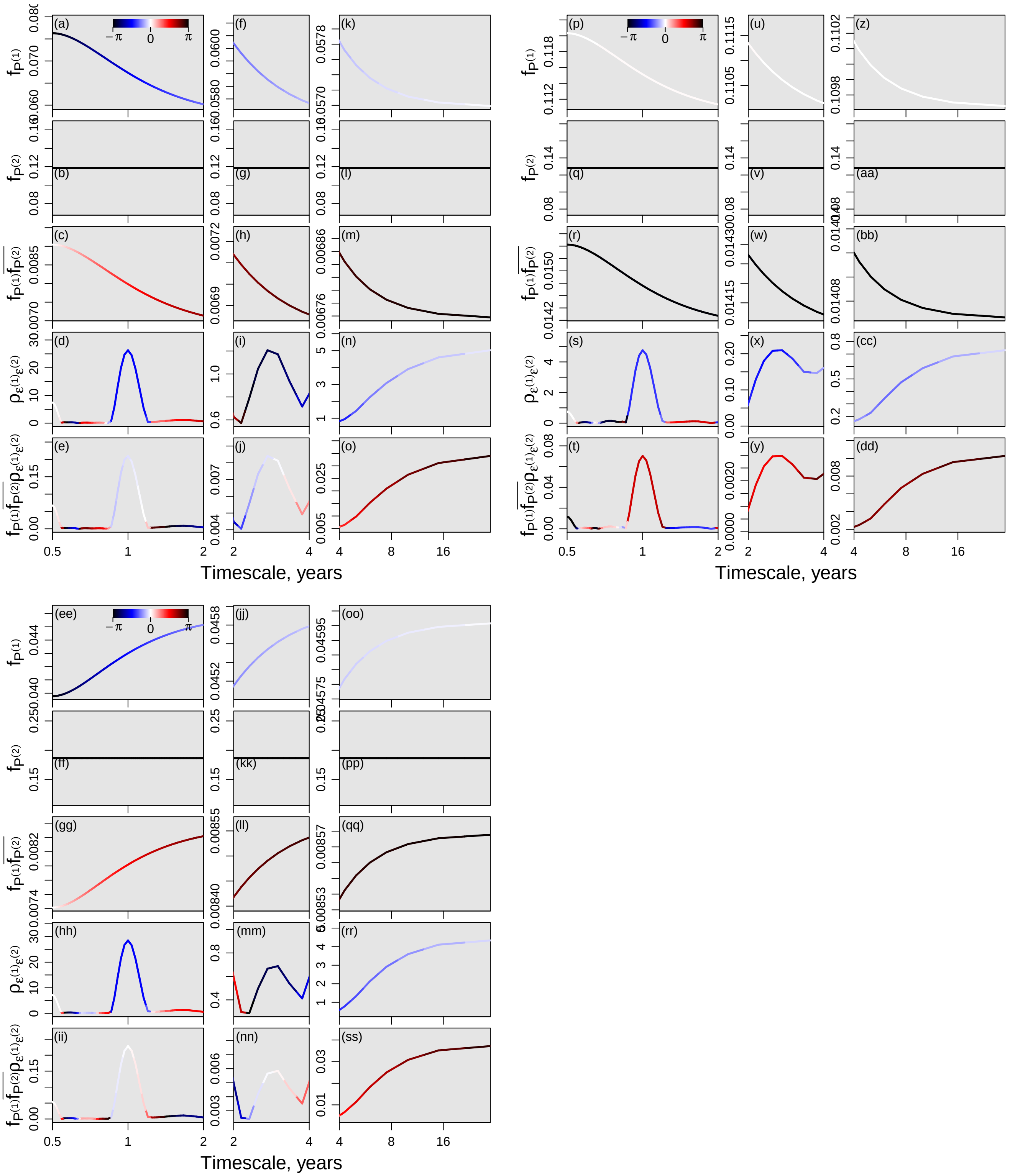

Figure S10: Same format as Fig. S8, but based on a model that considered potential kelp lag effects from up to 12 quarters in the past (see Methods). Results were substantively the same as for Fig. S8.

Table S1: Regression coefficients used to parameterize (1)-(4) of the main text for each of our three regions. All values rounded to two decimal places.

| | $p_0^{(1)}$ | $p_1^{(1)}$ | $p_0^{(2)}$ | $b_1, b_2, \ldots, b_n$ | $R^2$ |
|---|---|---|---|---|---|
| CCA1, $n = 4$ | -0.03 | 0.11 | -0.23 | 0.26,-0.22,0.11,0.21 | 0.57 |
| CCA2, $n = 4$ | 0 | 0.07 | -0.24 | 0.31,-0.14,0.09,0.16 | 0.38 |
| SB, $n = 4$ | 0.13 | -0.02 | -0.12 | 0.31,-0.03,-0.07,0.17 | 0.19 |
| CCA1, $n = 8$ | -0.02 | 0.08 | -0.21 | 0.36,-0.22,0.09,0.24,-0.23,0.02,-0.1,0.09 | 0.6 |
| CCA2, $n = 8$ | 0 | 0.06 | -0.22 | 0.37,-0.14,0.08,0.17,-0.14,0.01,0,0.1 | 0.4 |
| SB, $n = 8$ | 0.13 | -0.02 | -0.13 | 0.32,-0.04,-0.05,0.17,-0.07,0.03,-0.04,0.06 | 0.2 |
| CCA1, $n = 12$ | -0.01 | 0.07 | -0.12 | 0.35,-0.19,0.09,0.17,-0.16,0.04,-0.09,0.09,-0.12,-0.05,-0.06,0.2 | 0.63 |
| CCA2, $n = 12$ | 0 | 0.04 | -0.19 | 0.39,-0.13,0.08,0.13,-0.1,0.03,-0.03,0.14,-0.14,-0.03,-0.01,0.11 | 0.43 |
| SB, $n = 12$ | 0.12 | -0.01 | -0.13 | 0.33,-0.03,-0.06,0.15,-0.06,0.02,-0.04,0.08,-0.09,-0.03,0,0.08 | 0.22 |
